# Injectable ventral spinal stimulator evokes programmable and biomimetic hindlimb motion

**DOI:** 10.1101/2023.06.15.545178

**Authors:** Dingchang Lin, Jung Min Lee, Chonghe Wang, Hong-Gyu Park, Charles M. Lieber

## Abstract

Spinal cord neuromodulation can restore partial to complete loss of motor functions associated with neuromotor disease and trauma. Current technologies have made substantial progress, but have limitations as dorsal epidural or intraspinal devices that are either remote to ventral motor neurons or subject to surgical intervention in the spinal tissue. Here, we describe a flexible and stretchable spinal stimulator design with nanoscale thickness that can be implanted by minimally-invasive injection through a polymeric catheter to target the ventral spinal space of mice. Ventrolaterally implanted devices exhibited substantially lower stimulation threshold currents and more precise recruitment of motor pools than comparable dorsal epidural implants. Functionally relevant and novel hindlimb movements were achieved via specific stimulation patterns of the electrodes. This approach holds translational potential for improving controllable limb function following spinal cord injury or neuromotor disease.

---

The spinal cord is a key element of the central nervous system (CNS) for the transmission of descending motor commands, ascending sensory inputs, and producing adaptive motor patterns ^1^. Spinal cord injury (SCI) and other neuromotor diseases can disrupt these pathways and circuits and eliminate sensory and motor function below the trauma ^1, 2^.

For paralyzed individuals, restoring basic leg motor functions represents a high priority. A myriad of efforts have been pursued to develop bio-integrated electronics to achieve this goal ^3-6^. For example, functional electrical stimulation (FES) via peripheral nerves or muscles holds promise and has been extensively employed in rehabilitation and physical therapy ^5, 6^. When applied to peripheral nerves or muscles, FES elicits a large force to support the body weight; and by alternating the recruitment of flexors and extensors, command-driven walking can be fulfilled with the aid of crutches or walkers. Nevertheless, rapid muscle fatigue and difficulties in controlling individual muscles in FES lead to a restricted functional walking distance that is less applicable for daily-life recovery of leg functions after paralysis ^5, 7^.

Spinal cord stimulators are a promising alternative to restore at least partial motor control following SCI ^8-12^. In particular, the compact device size, resistance to fatigue during continuous stimulation, and low power consumption of spinal cord stimulation (SCS) offer the potential to restore motor control in daily life, especially long-distance walking. In addition, SCS can facilitate recovery from SCI by promoting axon regeneration ^13, 14^, driving neuroplastic change of spinal circuits ^15, 16^, and maintaining high basal excitability of motoneurons ^17-19^. Hence, SCS can both provide immediate limb motor control and also facilitate long-term motor and sensory recovery.

Neuroprosthetic devices for dorsal epidural electrical stimulation (dEES) ^20-22^ and intraspinal microstimulation (ISMS) ^23-25^ have shown promising results but also have limitations ^11^. On one hand, dEES devices exhibit low surgical lesions and long-term biocompatibility, yet the electrodes are distal to the ventral motor pools, and thus leg movements are primarily elicited by transsynaptic coupling that enhances basal neuronal excitability ^19^. Therefore, serotonergic replacement therapy as well as residual descending and proprioceptive sensory inputs are often needed ^17, 26^. On the other hand, ISMS devices can directly activate motoneurons and the fibers-in-passage to elicit functional movements with high precision and fidelity, but these intraspinal devices entail greater surgical complexity, poor chronic stability, and can damage spinal tracts ^11^. To date, methods that simultaneously exhibit minimally invasive surgical procedures, stability, and efficient direct activation of motoneurons and fibers-in-passage, have been unavailable.

Anatomically, the ventrolateral epidural space is closest to the motoneurons without penetrating the spinal tissue ^27^, and thus we hypothesized that implanting SCS devices that are sufficiently small, flexible and stretchable in this region could boost the efficacy and precision of activation while preserving the low invasiveness and chronic compatibility of dEES ^28^. Previously, this strategy has been rarely taken ^29^, and we speculated that this was due to the low accessibility of the ventral spinal cord with reported epidural devices ^10, 22^. In particular, existing dEES devices adopt the same design and technologies for chronic pain alleviation ^20, 21^, which exhibit similar widths to the spinal cord and are thus unable to circumvent the laterally exited spinal nerves in posterior surgical approaches ^30^.

Here, we report a new design spinal cord stimulator that is miniaturized with nanoscale thickness and possesses ultraflexibility and stretchability necessary to be seamlessly implanted on the ventrolateral epidural surfaces of the mouse spinal cord using a simple syringe injection through flexible polymer catheters. The implantation surgery and newly designed SCS devices exhibit negligible invasiveness with relatively rapid recovery of full motor function in mice. Ventrolateral EES (vlEES) shows over one order of magnitude lower threshold current, more diverse stimulated modes of motions than dEES as well as resistance to fatigue. Hindlimb movements with functional relevance, including bipedal motion, can be elicited by programmed vlEES via the multiple independently addressable electrodes in our new design devices.

We target the ventrolateral regions of the mouse spinal cord with an ultra-flexible and stretchable stimulator (**Fig. 1a**) injected via a flexible microcatheter into the epidural spinal space without the need to surgically expose the regions where electrodes are positioned (**Fig. 1b**). Briefly, the device preloaded in a microcatheter is implanted by ejecting a microliter volume of sterile saline, while the microcatheter is withdrawn (**Fig. 1c**). Ventrolateral access is readily achieved via insertion through 10^th^ and 11^th^ thoracic vertebrae (T10/T11) gap, where the concave local structure of rodent spine naturally guides the microcatheter to the ventral epidural spaceswithout the need for laminectomy. The SCS device, which we term ventral spinal cord electronics (VentrE), has design features to meet requirements of implantation and stable chronic stimulation: (1) a width compatible with loading and smooth ejection from the microcatheter; (2) sufficient flexibility and stretchability to survive bending associated with animal activities; and (3) orientation independence of the electrodes. This latter point recognizes that either surface of the VentrE yields optimal stimulation given that syringe implantation does not control up/down orientation.

**Fig. 1.**
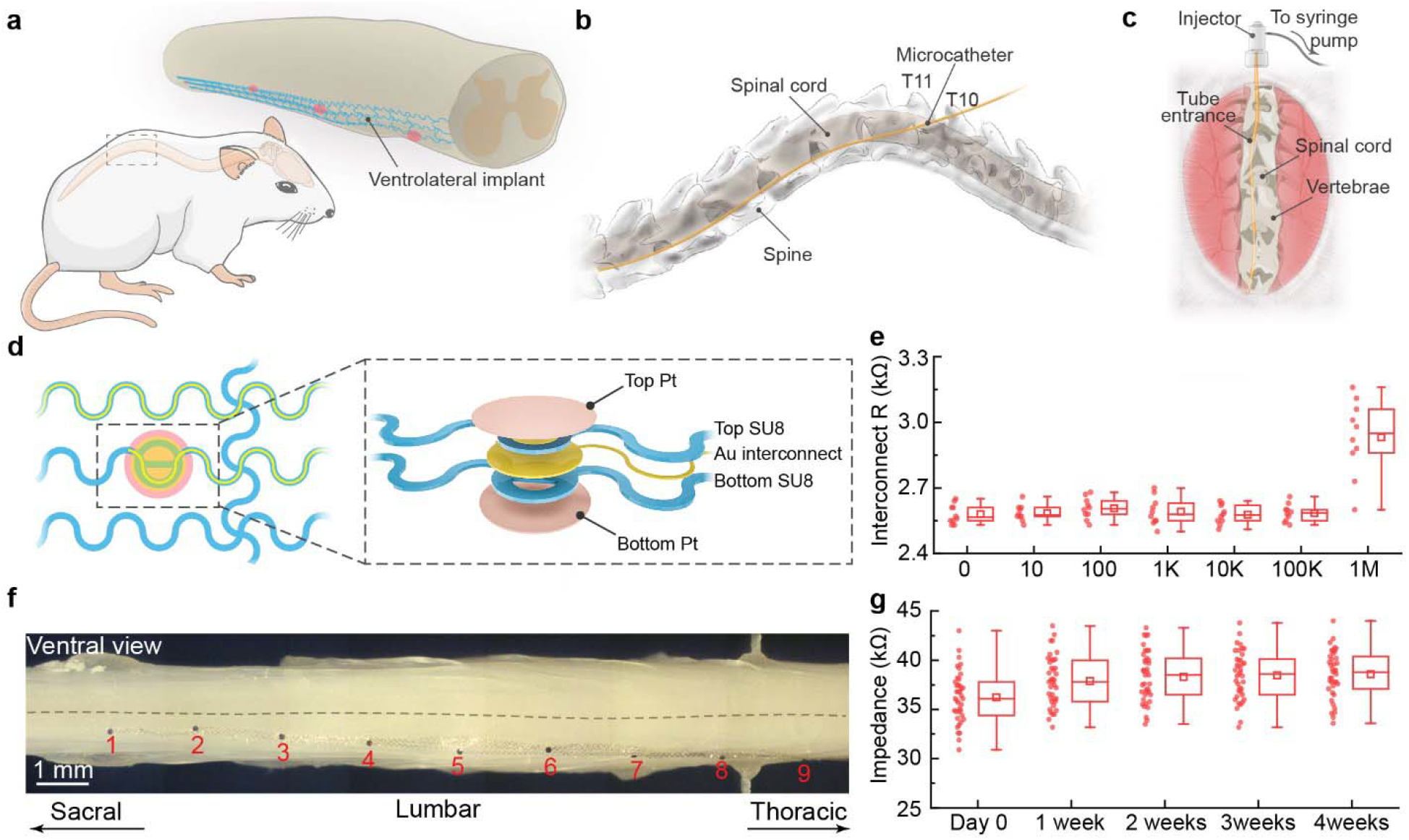
Syringe-injectable spinal stimulator. **a**, Schematic illustration of the spinal stimulator deployed on the ventrolateral epidural spaces. **b**, Schematic illustration of the method for delivering the stimulator to the ventrolateral surface of the lumbosacral segment. The microcatheter tube is inserted via the T10/T11 gap to access the ventrolateral epidural space. **c**, Schematic of syringe implantation of the device on the spinal cord epidural surface without laminectomy. **d**, Design of the stimulator backbone. Serpentine device ribbons were used to accommodate strains associated with spinal cord bending during normal animal activities. Schematic (right dash box) showing the layered structure of a double-sided stimulator microelectrode. **e**, Resistance evolution of the serpentine Au interconnects after uniaxial tensile fatigue cycles at a maximum strain of *ca*. 18 %. **f**, Stitched optical images showing the ventral epidural surface of the spinal cord with an implanted stimulator. The electrodes (*1 to 9*) are marked in sequence. **g**, *In vivo* evaluation of the implanted stimulator electrode impedances (at 1 kHz) up to 1 month post-surgery. Error bars: min-max; Box edges: 25% and 75%; Squares: mean; Lines: median.

VentrE design features that address these constraints are highlighted in **Fig. 1d, Fig. S1, and S2**. Overall, the devices are fabricated with a submicron-thick polymeric skeleton to yield a tissue-like bending stiffness described previously ^31-34^. To accommodate the surface strain of the bending spinal cord, a serpentine structure was incorporated in the longitudinal and transverse axes (**Fig. 1d *left*, Fig. S1**), allowing for stretching and compression without failure ^35^. Last, stimulation electrodes have a “double-sided” structure ^36^ such that a bare electrode surface contacts the spinal cord dura independent of implantation orientation (**Fig. 1d *right*, Fig. S2B**).

The serpentine design was guided by an estimated maximum surface strain of *ca*. 18% for the lumbosacral segment in adult mice (**Fig. S3, see methods**). Finite element analysis (FEA, **Fig. S4**) predicts that a structure with a semicircle periodicity of 100 μm and ribbon width of 10 μm (**Fig. 1b**) can sustain at least 30% strain without material failure. This estimate was validated by *in vitro* stretch and fatigue tests on the serpentine structure (**Fig. S5**). Indeed, tensile fatigue tests further demonstrated that the serpentine design can survive at least one million stretch cycles at *ca*. 18% strain, which was the maximum observed for freely-behaving mice, without breaking electrical connections (**Fig. 1e and Fig. S6**). Moreover, the designed structures remained mostly intact for up to 34.5% tensile strain (**Fig. S6**). These tests provide confidence that the VentrE design will yield robust chronic SCS in mice.

Ventrolateral implantation was carried out with the flexible microcatheter (OD: *ca*. 240 μm, ID: *ca*. 200 μm, **Fig. S7**) containing the VentrE inserted 1.6-1.8 cm along the mouse spinal canal via the T10/T11 gap without laminectomy (**Fig. 1b, c, Fig. S8**) allowing coverage of the major lumbosacral segment for hindlimb control ^30^. It is noteworthy that the dorsal epidural space can also be targeted using this microcatheter insertion, syringe-injection modality, by simply inserting the microcatheter via a lower thoracic gap (T11/T12, **Fig. S9**). The ventrolateral/dorsal position selectivity was confirmed by optical imaging of mouse spinal tissue from either ventrolateral (**Fig. 1f**) or dorsal (**Fig. S9C**) 4 weeks post-implantation at which time the serpentine metal interconnects were also intact based on optical microscopy analyses.

Given the structural stability inferred from the optical observations, we asked whether the stimulation microelectrodes remained electrically connected. This was evaluated by monitoring the impedance of implanted electrodes (N = 39 electrodes from 3 mice) weekly for four weeks post-implantation. During the 4-week evaluation period, the impedance of the electrodes remained stable in the range of *ca*. 30–45 kΩ without substantial time-dependent changes (**Fig. 1g**). The *in-vivo* evaluations corroborated the *in-vitro* tensile tests, confirming that the implanted stimulator array meets the stringent demands on mechanical properties for a spinal interface.

We then examined the invasiveness and long-term biocompatibility of implanted VentrE by comparing groups with VentrE implanted on both the dorsal and ventral spinal surfaces of each subject with a sham-operated group consisting of surgery and microcatheter insertion but no implantation. We first evaluated the overall motor activity post-surgery in an open field test (**Fig. S10, Movie S2**), where the traveling distance (**Fig. 2a**), average speed (**Fig. 2b**), and maximum speed (**Fig. 2c**) were quantified ^37^. The statistical analyses of the total traveling distance of both groups stabilized starting on day 2 and day 1, respectively (**Fig. 2a**), while the average (**Fig. 2b**) and the maximum speed (**Fig. 2c**) both stabilized on day 2 for both groups. A third group of mice without surgical operations (non-surgery group) were further used for comparison. The traveling distance, average speed, and maximum speed of both VentrE-implanted and sham-operated groups are indifferent from those of the non-surgery group following day 2. Therefore, we concluded that the mice in the VentrE-implanted group quickly recovered from the surgery and the implanted VentrE did not impact the long-term activity of the mice.

**Fig. 2.**
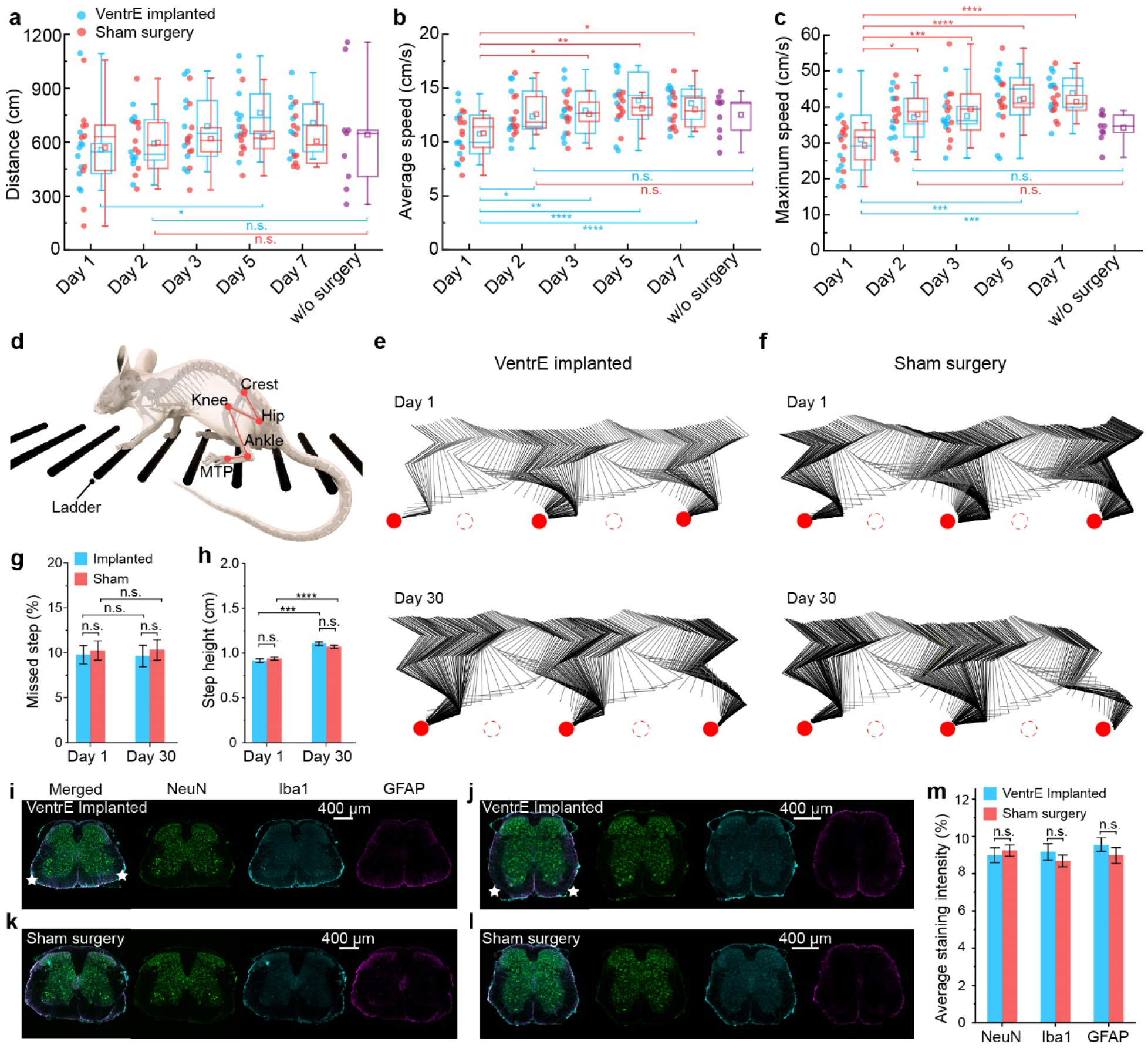
Evaluation of invasiveness and biocompatibility. **a-c**, Box plots comparing the activity as mean distance (**a**), average speed (**b**), and maximum speed (**c**) of free-moving mice with implanted VentrE (*blue*), sham surgeries (*red*), and without surgery (*violet*). (N = 5 mice per group, 2 trials per animal for sham-operated and stimulator-implanted surgeries; N = 9 mice, 1 trial per animal for the non-surgery group). Error bars: min-max; Box edges: 25% and 75%; Squares: mean; Lines: median. * P<0.05, ** P< 0.01; *** P< 0.001; **** P<0.0001. **d**, Schematic of the horizontal ladder rung walking test. The iliac crest, hip, knee, ankle, and metatarsophalangeal (MTP) joints are labeled and recognized for kinematic analyses. **e-f**, Hindlimb kinematics of a mouse with VentrE-implanted (**e**) and sham-surgery (**f**) during ladder walking 1 day (*top*) and 30 days (*bottom*) post-surgery. The red solid and dash circles indicate the positions of the rungs that the tracked and the contralateral limbs stepped on, respectively. **g**, Bar plots comparing the mean percentage of missed steps on rungs of the ladder between mice with stimulator-implanted (*blue*) and sham surgery (*red*) (N = 6 mice per group, 3 trials per animal). **h**, Bar plots comparing the mean step height between mice with VentrE-implanted (*blue*) and sham surgeries (*red*) (N = 6 mice per group, 1 trial per animal). **i-l**, Fluorescence images of the representative lumbar (**i** and **k**) and sacral (**j** and **l**) coronal sections with VentrE-implanted (**i** and **j**) and sham-surgery (**k** and **l**). The mice were sacrificed 2 weeks after surgery. Neurons (NeuN, green), microglia (Iba1, cyan), and astrocytes (GFAP, glial fibrillary acidic protein, magenta) are labeled in each slice. Stars indicate the approximate locations of implanted VentrEs. **m**, Statistic comparisons on the average staining intensity between VentrE-implanted and sham surgery groups (N = 3 mice per group, 4 coronal slices per mouse). Error bars: SEM. Statistical test: One-way and repeated-measure analysis of variance.

We then asked how this surgical procedure and VentE implantation affect skilled locomotor function by quantifying the hindlimb kinematics in a horizontal ladder rung walking test (**Fig. 2d, Fig. S11, Movie S3**) ^18, 38^. The tests were conducted on days 1 and 30 post-implantation to assess the immediate impact of the implantation procedures and the chronic biocompatibility, respectively. On day 1, the two groups showed similar hindlimb locomotion (**Fig. 2e, f** *top*, **Fig. S12A, B**). Statistical comparisons of their missed steps (**Fig. 2g**) and step height (**Fig. 2h**) did not show significant differences, indicating that the implantation is minimally invasive and does not compromise the hindlimb skilled locomotion. Furthermore, the two groups still exhibited comparable kinematic behavior (**Fig. 2e, f** *bottom*, **Fig. S12C, D**), percentage of missed steps (**Fig. 2g**), and step heights on day 30 (**Fig. 2h**), confirming the chronic stability of VentrE on the ventrolateral and dorsal spinal surfaces. It is noted that the step heights measured on day 30 were consistently higher than that on day 1 in both groups, which can be attributed to the gradual recovery of mice’s physical conditions after surgical operations. Comparison to reports for mice without surgery showed that the day-30 VentrE implanted and sham surgery mice had similar hindlimb locomotion ^18, 38^.

In addition, potential inflammatory responses were evaluated at the tissue and cellular levels, by quantifying the distributions of neurons, astrocytes, and microglia in the coronal sections of the lumbosacral segments (**Fig. 2i-m, Fig. S13**) ^18^. Neither visual observation (**Fig. 2i-l, Fig. S13, S14A-D**) nor quantification (**Fig. 2m, Fig. S14E**) of neuron distributions in the grey matter revealed significant differences between the two groups. Comparison of astrocytes (anti-gfap) and the microglia (anti-Iba1), also reveals similar distributions at the cellular level between the implanted and sham-operated groups at both two (**Fig. 2i-m, Fig. S13**) and four weeks (**Fig. S14**) post-surgery. The results indicate negligible neuronal depletion and inflammatory responses at the local spinal cord tissue after VentrE implantation and are consistent with minimal invasiveness for the implanted VentrE.

We next asked how the basic characteristics of vlEES versus dEES of hindlimb movements compared for implanted VentrE devices in anesthetized mice. This initial phase of our VentrE stimulation studies focused on motor thresholds, evoked motion modes, and fatigue to prolonged stimulation with the anesthetized mice in the suspended configuration (**Fig. 3a**), as used previously in rat dorsal EES studies ^39^.

**Fig. 3.**
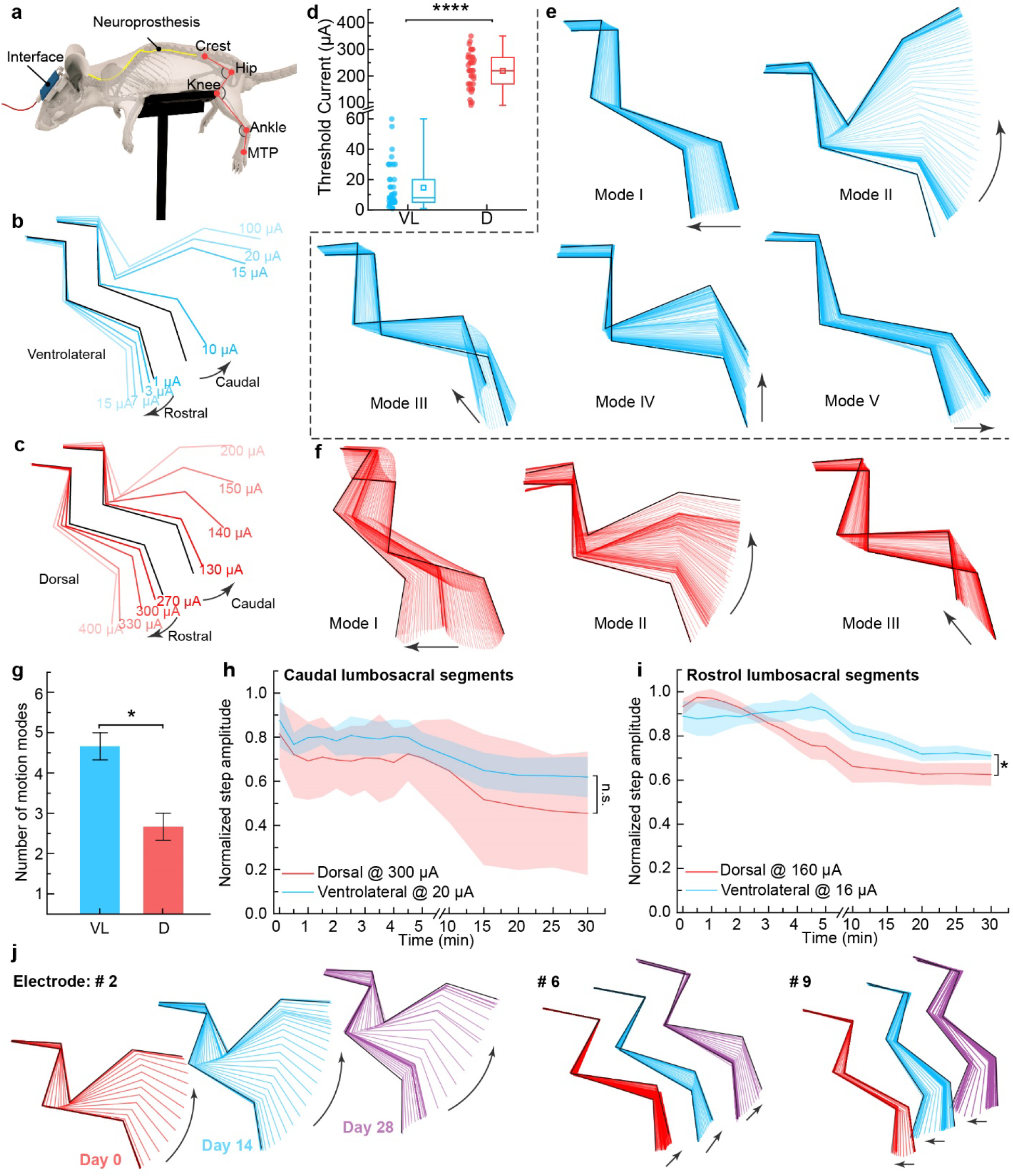
Ventrolateral vs. dorsal implant stimulation. **a**, Schematic of the suspended hindlimb kinematics. An anesthetized mouse is placed on the suspension stage with the iliac crest, hip, knee, ankle, and MTP joints labeled and recognized for offline analysis. The VentrE interface was mounted on the skull and connected during measurements to a computer-controlled stimulator. **b-c**, Representative hindlimb kinematics vs. stimulation currents delivered via vlEES (**b**) or dEES (**c**) to the rostral/caudal lumbosacral segments. (**d**) Statistic distributions of the threshold currents for electrodes on the ventrolateral (*blue*) and dorsal (*red*) implants (40 electrodes from N = 3 mice each). Error bars: min-max; Box edges: 25% and 75%; Squares: mean; Lines: median. **e-f**, Representative stimulation modes elicited by VentrE implanted to the ventrolateral (**E**) and dorsal (**f**) surfaces of the lumbosacral segments. (**g**) The number of motion modes activated by the ventrolateral (*blue*) and dorsal (*red*) implants (N = 3 mice each). **h-i**, Evolution of mean hindlimb step amplitudes versus time when stimulating via the ventrolateral (*blue*) and the dorsal (*red*) implants at the caudal (**h**) and the rostral (**i**) lumbosacral segments (N = 4 trials each condition). The edges of the shaded areas reflect the standard deviation (± S.D.). * P< 0.05; **** P<0.0001. Statistical test: One-way analysis of variance. **j**, Chronic studies of hindlimb kinematics showing the motion modes elicited by VentrE elecltrodes 2, 6 and 9 on days 0, 14 and 28.

Studies of vlEES and dEES (**Fig. 3b,c**; **Fig. S15**) showed that the mouse hindlimb responded to significantly lower currents for vlEES. Specifically, vlEES exhibited hindlimb movement onsets at < 10 μA and sometimes as low as 1 μA (**Fig. 3b**). In contrast, dEES required *ca*. 100 μA and sometimes over 250 μA to evoke hindlimb motion. The substantially lower threshold for vlEES was confirmed by statistical analysis (N = 40 electrodes from 3 mice per group) of the motor threshold, defined as the average onset current that evokes distinguishable hindlimb movements. We observed over one order of magnitude lower motor thresholds for vlEES vs. dEES, 14.7 ± 2.3 vs. 219.3 ± 10.7 (SEM) μA, respectively (**Fig. 3d**). It is noteworthy that 17.5% of the investigated ventrolateral electrodes have a threshold ≤ 5 μA, equivalent to an imposed charge in the picocoulomb (pC) range. The low imposed charge should minimize undesirable Faradic processes during stimulation that can adversely affect spinal tissue over time ^40^.

These stimulation threshold studies also showed that more diverse motor activity could be elicited by vlEES than dEES. Here, we exploited the hindlimb kinematics analyses ^39^ (**see methods**) to evaluate the possible modes of hindlimb motion elicited by individual electrodes, each of which represents a pattern of synergetic recruitment of the flexors and extensors in the mouse hindlimbs. Each motion mode was elicited by four pulses of 200 μs in a 40 Hz stimulation train delivered via an electrode on VentrE (Supporting Information). Specifically, five distinct modes of motion could be elicited using individual electrodes of a 9-channel stimulation array implanted on the epidural lumbosacral segment (**Fig. 3e**), while dEES using the same design VentrE typically showed two or three distinct modes (**Fig. 3f**). A statistical summary reveals that vlEES showed 4.7 ± 0.3 (SEM) modes or 75% more than that of dEES, 2.7 ± 0.3 (**Fig. 3g**). We suggest that the low threshold currents and more diverse motor modes found with our vlEES can be attributed to the close proximity of the stimulation sites to the motoneurons and the efferent fibers. This idea is supported by computations that showed a more concentrated electric field with a shorter distance between recruited components for EES ^41^, which can allow more efficient and precise recruitment of the ventral motoneurons.

The ventrolateral stimulation sites showed comparable or improved resistance to fatigue during continuous stimulation studies. The decay of hindlimb step amplitudes was recorded using a continuous stimulation protocol (**see Methods**) similar to that reported elsewhere ^42^ to electrodes on caudal and rostral spinal lumbosacral segments.

In general, dEES already exhibits good resistance to fatigue in both caudal (**Fig. 3h**) and rostral (**Fig. 3i**) lumbosacral sites, with an average amplitude retention of 45.5% and 62.6% after 30 min continuous stimulation, respectively. The results are consistent with previous studies that show good resistance to fatigue ^11, 42^. In comparison, vlEES exhibits slightly better resistance to fatigue, where average amplitude retention of 61.9% and 71.0% were obtained on the caudal and rostral sites, respectively (**Fig. 3h, i**). Since fast fatigue generally originated from the recruitments of large axons that innervate the more fatigable muscle fibers ^43^, the result illustrates that vlEES might preferentially engage motoneurons and fatigue-resistant fibers.

We further evaluated the capability of VentrE implants to evoke chronically-stable hindlimb motions, where hindlimb kinematics at day 0, 14, and 28 post-surgery were analyzed (**Fig. 3j**). The evoked motions at various times are comparable for each of the individual electrodes tested, indicating stable VentrE positioning on the ventrolateral epidural surface and recruitment of motor pools during stimulation. The results are also consistent with post measurement VentrE/spinal tissue analyses (**Fig. 1f, Fig. S9C**) that showed the VentrE implants adhered strongly to the epidural spinal surface. Taken together, these results illustrate that the ultraflexibility and stretchability of VentrE are crucial to ensure a minimally-invasive yet tight electrode/spinal tissue interface that provides chronically-stable hindlimb control.

The distinct results for ventrolateral stimulation led us to ask whether it would be possible to program stimulation patterns that evoke more complex and biomimetic hindlimb movements. By recruiting multiple electrodes in sequence (**See Methods**), we elicited functionally relevant hindlimb modes including pedaling (**Fig. 4a, Fig. S16A**), kicking (**Fig. 4b, Fig. S16B**), and waving (**Fig. 4c, Fig. S16C**) in anesthetized mice (**Movie S4**). In addition, we asked whether bilateral stimulation by two VentrEs implanted on the lumbosacral segment (**Fig. 4d**) could enable bipedal hindlimb motion. Multi-train stimulation applied to either rostral or caudal lumbosacral segments using the two implants (**See Methods**) elicited similar modes of waving (**Figs. 4e, f**). Significantly, simultaneously applying the left/right stimulation trains offset by 0.4 s to the bilateral VentrE implants (Methods) elicits bipedal locomotor-like activity (**Fig. 4g, Movie S5**), whereas ventrolateral stimulation with one or the other VentrE elicits only left or right pedal motion. The detailed kinematic progression (**Fig. 4g**) showed an alternating motion of the left (*top*) and the right (*bottom*) hindlimbs.

**Fig. 4.**
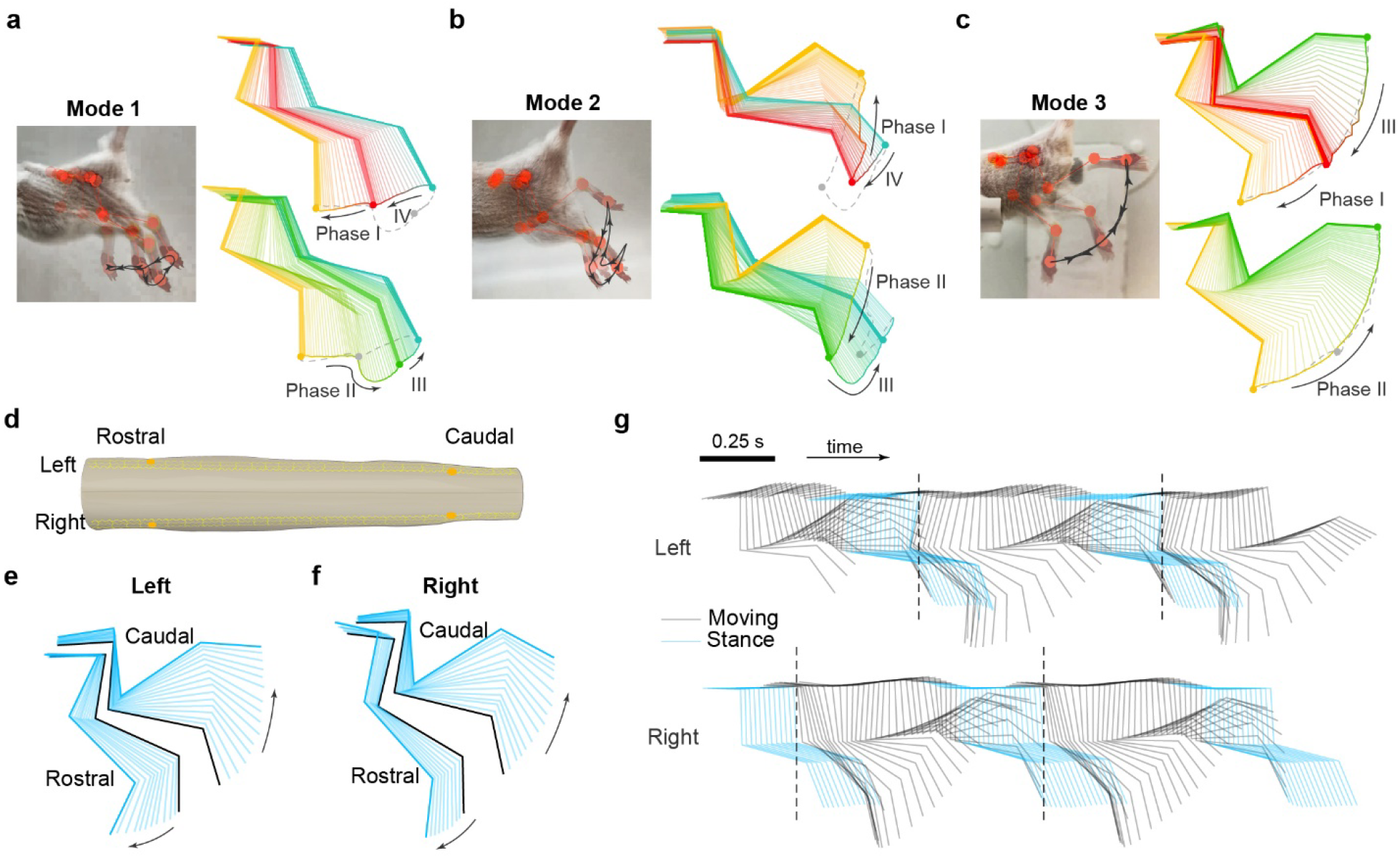
Unipedal and bipedal hindlimb activity. **a**-**c** Superimposed video sequences of the elicited motions in one cycle (left), and the corresponding kinematic analyses (right). Three functionally relevant motions corresponding to pedaling (**a**), kicking (**b**), and waving (**c**) were shown. Each programmed motion is segmented into distinct phases in the kinematic analyses. **d**. Schematic illustration of the bilateral implantation to lumbosacral spinal segments. The approximate positions of the electrodes employed for bipedal programmed stimulation were labeled accordingly. (**e** and **f**) Motion evoked by simultaneous stimulation on the left side (**e**) and the right (**f**) side of the lumbosacral spinal segment. The stimulation trains applied to the bilateral VentrE implants were similar and offset as described in the Methods. (**g**) The kinematic progression of the left (*top*) and the right (*bottom*) hindlimb during bipedal programmed stimulation. The onsets of motion are marked with dash lines.

These studies in mice show that the ventrolateral epidural spinal surface is a promising target for SCS. We demonstrated minimally invasive syringe implantation of ultra-flexible and stretchable neuroprosthetic devices that stimulate the spinal cord via the ventrolateral epidural space. The VentrE has excellent biocompatibility and resulting vlEES shows high stimulation efficacy and resistance to fatigue. The low threshold current and diverse modes of evoked motion are especially interesting compared to previous studies of dEES. In addition, these characteristics of the VentrE implants were used in vlEES to elicit functionally relevant hindlimb modes, including natural bipedal motion via timed pulse trains applied to bilateral VentrE implants. We suggest that the high stimulation efficacy and precision can be attributed to the close proximity of ventral epidural sites to motoneurons and fibers-in-passage compared to more remote dorsal stimulation. In the future, the VentrE electrode positioning for vlEES could be further optimized using real-time fluoroscopy imaging ^44^.

This study represents the beginning of work on and potential translation of vlEES where both fundamental and engineering studies will be necessary in the future. For example, it will be interesting to investigate the mechanism underlying the attractive characteristics of vlEES such as cellular-level recruitment and functional connectivity of spinal motor circuits ^45, 46^. Intramuscular electromyography (EMG) would also be useful to quantify detailed muscle activation by individual electrodes of the implanted VentrE in vlEES. Last, evaluation of the capability to restore weight-bearing locomotion in rodent models with SCI would be a starting point to explore for translation where ultimately nonhuman primates and humans, which have much larger spinal canals than mice, would be of interest.

## Supporting Information

The Supporting Information is available free of charge at https://nam02.safelinks.protection.outlook.com/?url=http%3A%2F%2Fpubs.acs.org%2F&data=05%7C01%7Cdclin%40jhu.edu%7C22500190bdbc4890d70708db6b6a7eff%7C9fa4f438b1e6473b803f86f8aedf0dec%7C0%7C0%7C638221877362060714%7CUnknown%7CTWFpbGZsb3d8eyJWIjoiMC4wLjAwMDAiLCJQIjoiV2luMzIiLCJBTiI6Ik1haWwiLCJXVCI6Mn0%3D%7C3000%7C%7C%7C&sdata=WTzSv62mDE7G9vkddv1Di45VHLqjr7VI8vDx1rZMkH8%3D&reserved=0.%E2%80%9D

Additional experimental data, including the detailed experimental methods, mechanical measurements and evaluation, probe design, animal behavioral tests, histology, and kinematic analyses (PDF).

## ACKNOWLEDGMENTS

We thank Dr. Guosong Hong, Dr. Tao Zhou, Yitong Qi, and Dr. Yayuan Liu for their helpful discussions. D.L. acknowledges the support of this work by the Johns Hopkins University Start-up Fund. C.M.L. thanks the Lieber Research Group for their strong support. This work was carried out in part at the Singh Center for Nanotechnology, part of the National Nanotechnology Coordinated Infrastructure Program, which is supported by the National Science Foundation grant NNCI-2025608. H.-G.P. acknowledges support from a National Research Foundation of Korea (NRF) grant funded by the Korean government (MSIT) (2021R1A2C3006781).

## Author contributions

D.L. and C.M.L. conceived the concept and designed the experiments; D.L., and J.M.L. performed the experiments; D.L. wrote Python codes for data analyses. D.L., J.M.L., and C.M.L. analyzed the data and wrote the paper; all authors discussed the results and commented on the manuscript.

## Competing interests

None.

## Data and materials availability

All data are available in the manuscript or the supplementary materials. Devices and experimental materials are available upon request to the authors.

## Notes

### Competing Interest Statement

The authors have declared no competing interest.

## REFERENCES AND NOTES

1. Kiehn, O. Decoding the organization of spinal circuits that control locomotion. Nature Reviews Neuroscience 2016, 17, 224.

2. Damasio, A.; Carvalho, G. B. The nature of feelings: evolutionary and neurobiological origins. Nature Reviews Neuroscience 2013, 14, (2), 143–152.

3. Rodríguez-Fernández, A.; Lobo-Prat, J.; Font-Llagunes, J. M. Systematic review on wearable lower-limb exoskeletons for gait training in neuromuscular impairments. Journal of NeuroEngineering and Rehabilitation 2021, 18, (1), 22.

4. Dupont, P. E.; Nelson, B. J.; Goldfarb, M.; Hannaford, B.; Menciassi, A.; O’Malley, M. K.; Simaan, N.; Valdastri, P.; Yang, G.-Z. A decade retrospective of medical robotics research from 2010 to 2020. Science Robotics 2021, 6, (60), eabi8017.

5. Marquez-Chin, C.; Popovic, M. R. Functional electrical stimulation therapy for restoration of motor function after spinal cord injury and stroke: a review. BioMedical Engineering OnLine 2020, 19, (1), 34.

6. Mushahwar, V. K.; Jacobs, P. L.; Normann, R. A.; Triolo, R. J.; Kleitman, N. New functional electrical stimulation approaches to standing and walking. Journal of Neural Engineering 2007, 4, (3), S181–S197.

7. Fouad, K.; Tetzlaff, W. Rehabilitative training and plasticity following spinal cord injury. Experimental neurology 2012, 235, (1), 91–99.

8. Fuentes, R.; Petersson, P.; Siesser, W. B.; Caron, M. G.; Nicolelis, M. A. L. Spinal Cord Stimulation Restores Locomotion in Animal Models of Parkinson’s Disease. Science 2009, 323, (5921), 1578–1582.

9. Jackson, A.; Zimmermann, J. B. Neural interfaces for the brain and spinal cord—restoring motor function. Nature Reviews Neurology 2012, 8, (12), 690–699.

10. Borton, D.; Micera, S.; Millán, J. d. R.; Courtine, G. Personalized Neuroprosthetics. Science Translational Medicine 2013, 5, (210), 210rv2–210rv2.

11. Ievins, A.; Moritz, C. T. Therapeutic Stimulation for Restoration of Function After Spinal Cord Injury. Physiology 2017, 32, (5), 391–398.

12. Taccola, G.; Sayenko, D.; Gad, P.; Gerasimenko, Y.; Edgerton, V. R. And yet it moves: Recovery of volitional control after spinal cord injury. Progress in Neurobiology 2018, 160, 64–81.

13. Chen, B.; Li, Y.; Yu, B.; Zhang, Z.; Brommer, B.; Williams, P. R.; Liu, Y.; Hegarty, S. V.; Zhou, S.; Zhu, J.; Guo, H.; Lu, Y.; Zhang, Y.; Gu, X.; He, Z. Reactivation of Dormant Relay Pathways in Injured Spinal Cord by KCC2 Manipulations. Cell 2018, 174, (3), 521-535.e13.

14. Hamid, S.; Hayek, R. Role of electrical stimulation for rehabilitation and regeneration after spinal cord injury: an overview. European Spine Journal 2008, 17, (9), 1256–1269.

15. and, J. R. W.; Tennissen, A. M. Activity-Dependent Spinal Cord Plasticity in Health and Disease. Annual Review of Neuroscience 2001, 24, (1), 807–843.

16. Carhart, M. R.; He, J.; Herman, R.; D’luzansky, S.; Willis, W. T. Epidural spinal-cord stimulation facilitates recovery of functional walking following incomplete spinal-cord injury. IEEE Transactions on neural systems and rehabilitation engineering 2004, 12, (1), 32–42.

17. Courtine, G.; Gerasimenko, Y.; van den Brand, R.; Yew, A.; Musienko, P.; Zhong, H.; Song, B.; Ao, Y.; Ichiyama, R. M.; Lavrov, I.; Roy, R. R.; Sofroniew, M. V.; Edgerton, V. R. Transformation of nonfunctional spinal circuits into functional states after the loss of brain input. Nature Neuroscience 2009, 12, (10), 1333–1342.

18. Minev, I. R.; Musienko, P.; Hirsch, A.; Barraud, Q.; Wenger, N.; Moraud, E. M.; Gandar, J.; Capogrosso, M.; Milekovic, T.; Asboth, L.; Torres, R. F.; Vachicouras, N.; Liu, Q.; Pavlova, N.; Duis, S.; Larmagnac, A.; Vörös, J.; Micera, S.; Suo, Z.; Courtine, G.; Lacour, S. P. Electronic dura mater for longterm multimodal neural interfaces. Science 2015, 347, (6218), 159–163.

19. Angeli, C. A.; Edgerton, V. R.; Gerasimenko, Y. P.; Harkema, S. J. Altering spinal cord excitability enables voluntary movements after chronic complete paralysis in humans. Brain 2014, 137, (5), 1394–1409.

20. Wagner, F. B.; Mignardot, J.-B.; Le Goff-Mignardot, C. G.; Demesmaeker, R.; Komi, S.; Capogrosso, M.; Rowald, A.; Seáñez, I.; Caban, M.; Pirondini, E.; Vat, M.; McCracken, L. A.; Heimgartner, R.; Fodor, I.; Watrin, A.; Seguin, P.; Paoles, E.; Van Den Keybus, K.; Eberle, G.; Schurch, B.; Pralong, E.; Becce, F.; Prior, J.; Buse, N.; Buschman, R.; Neufeld, E.; Kuster, N.; Carda, S.; von Zitzewitz, J.; Delattre, V.; Denison, T.; Lambert, H.; Minassian, K.; Bloch, J.; Courtine, G. Targeted neurotechnology restores walking in humans with spinal cord injury. Nature 2018, 563, (7729), 65–71.

21. Gill, M. L.; Grahn, P. J.; Calvert, J. S.; Linde, M. B.; Lavrov, I. A.; Strommen, J. A.; Beck, L. A.; Sayenko, D. G.; Van Straaten, M. G.; Drubach, D. I.; Veith, D. D.; Thoreson, A. R.; Lopez, C.; Gerasimenko, Y. P.; Edgerton, V. R.; Lee, K. H.; Zhao, K. D. Neuromodulation of lumbosacral spinal networks enables independent stepping after complete paraplegia. Nature Medicine 2018, 24, (11), 1677–1682.

22. Rowald, A.; Komi, S.; Demesmaeker, R.; Baaklini, E.; Hernandez-Charpak, S. D.; Paoles, E.; Montanaro, H.; Cassara, A.; Becce, F.; Lloyd, B.; Newton, T.; Ravier, J.; Kinany, N.; D’Ercole, M.; Paley, A.; Hankov, N.; Varescon, C.; McCracken, L.; Vat, M.; Caban, M.; Watrin, A.; Jacquet, C.; Bole-Feysot, L.; Harte, C.; Lorach, H.; Galvez, A.; Tschopp, M.; Herrmann, N.; Wacker, M.; Geernaert, L.; Fodor, I.; Radevich, V.; Van Den Keybus, K.; Eberle, G.; Pralong, E.; Roulet, M.; Ledoux, J.-B.; Fornari, E.; Mandija, S.; Mattera, L.; Martuzzi, R.; Nazarian, B.; Benkler, S.; Callegari, S.; Greiner, N.; Fuhrer, B.; Froeling, M.; Buse, N.; Denison, T.; Buschman, R.; Wende, C.; Ganty, D.; Bakker, J.; Delattre, V.; Lambert, H.; Minassian, K.; van den Berg, C. A. T.; Kavounoudias, A.; Micera, S.; Van De Ville, D.; Barraud, Q.; Kurt, E.; Kuster, N.; Neufeld, E.; Capogrosso, M.; Asboth, L.; Wagner, F. B.; Bloch, J.; Courtine, G. Activity-dependent spinal cord neuromodulation rapidly restores trunk and leg motor functions after complete paralysis. Nature Medicine 2022, 28, (2), 260–271.

23. Dalrymple, A. N.; Roszko, D. A.; Sutton, R. S.; Mushahwar, V. K. Pavlovian control of intraspinal microstimulation to produce over-ground walking. Journal of Neural Engineering 2020, 17, (3), 036002.

24. Moritz, C. T.; Lucas, T. H.; Perlmutter, S. I.; Fetz, E. E. Forelimb Movements and Muscle Responses Evoked by Microstimulation of Cervical Spinal Cord in Sedated Monkeys. Journal of Neurophysiology 2007, 97, (1), 110–120.

25. Sunshine, M. D.; Cho, F. S.; Lockwood, D. R.; Fechko, A. S.; Kasten, M. R.; Moritz, C. T. Cervical intraspinal microstimulation evokes robust forelimb movements before and after injury. Journal of Neural Engineering 2013, 10, (3), 036001.

26. Fouad, K.; Rank, M. M.; Vavrek, R.; Murray, K. C.; Sanelli, L.; Bennett, D. J. Locomotion After Spinal Cord Injury Depends on Constitutive Activity in Serotonin Receptors. Journal of Neurophysiology 2010, 104, (6), 2975–2984.

27. Watson, C.; Paxinos, G.; Kayalioglu, G.; Heise, C., Chapter 16 - Atlas of the Mouse Spinal Cord. In The Spinal Cord, Watson, C.; Paxinos, G.; Kayalioglu, G., Eds. Academic Press: San Diego, 2009; pp 308–379.

28. Sharpe, A. N.; Jackson, A. Upper-limb muscle responses to epidural, subdural and intraspinal stimulation of the cervical spinal cord. Journal of Neural Engineering 2014, 11, (1), 016005.

29. Hogan, M. K.; Barber, S. M.; Rao, Z.; Kondiles, B. R.; Huang, M.; Steele, W. J.; Yu, C.; Horner, P. J. A wireless spinal stimulation system for ventral activation of the rat cervical spinal cord. Scientific Reports 2021, 11, (1), 14900.

30. Harrison, M.; O’Brien, A.; Adams, L.; Cowin, G.; Ruitenberg, M. J.; Sengul, G.; Watson, C. Vertebral landmarks for the identification of spinal cord segments in the mouse. NeuroImage 2013, 68, 22–29.

31. Fu, T.-M.; Hong, G.; Zhou, T.; Schuhmann, T. G.; Viveros, R. D.; Lieber, C. M. Stable long-term chronic brain mapping at the single-neuron level. Nature Methods 2016, 13, 875.

32. Hong, G.; Fu, T.-M.; Qiao, M.; Viveros, R. D.; Yang, X.; Zhou, T.; Lee, J. M.; Park, H.-G.; Sanes, J. R.; Lieber, C. M. A method for single-neuron chronic recording from the retina in awake mice. Science 2018, 360, (6396), 1447–1451.

33. Yang, X.; Zhou, T.; Zwang, T. J.; Hong, G.; Zhao, Y.; Viveros, R. D.; Fu, T.-M.; Gao, T.; Lieber, C. M. Bioinspired neuron-like electronics. Nature Materials 2019, 18, (5), 510–517.

34. Lee, J. M.; Hong, G.; Lin, D.; Schuhmann, T. G.; Sullivan, A. T.; Viveros, R. D.; Park, H.-G.; Lieber, C. M. Nanoenabled Direct Contact Interfacing of Syringe-Injectable Mesh Electronics. Nano Letters 2019, 19, (8), 5818–5826.

35. Zhang, Y.; Wang, S.; Li, X.; Fan, J. A.; Xu, S.; Song, Y. M.; Choi, K.-J.; Yeo, W.-H.; Lee, W.; Nazaar, S. N.; Lu, B.; Yin, L.; Hwang, K.-C.; Rogers, J. A.; Huang, Y. Experimental and Theoretical Studies of Serpentine Microstructures Bonded To Prestrained Elastomers for Stretchable Electronics. Advanced Functional Materials 2014, 24, (14), 2028–2037.

36. Lee, J. M.; Lin, D.; Hong, G.; Kim, K.-H.; Park, H.-G.; Lieber, C. M. Scalable Three-Dimensional Recording Electrodes for Probing Biological Tissues. Nano Letters 2022, 22, (11), 4552–4559.

37. Kraeuter, A.-K.; Guest, P. C.; Sarnyai, Z., The Open Field Test for Measuring Locomotor Activity and Anxiety-Like Behavior. In Pre-Clinical Models: Techniques and Protocols, Guest, P. C., Ed. Springer New York: New York, NY, 2019; pp 99–103.

38. Au-Metz, G. A.; Au-Whishaw, I. Q. The Ladder Rung Walking Task: A Scoring System and its Practical Application. JoVE 2009, (28), e1204.

39. Capogrosso, M.; Wagner, F. B.; Gandar, J.; Moraud, E. M.; Wenger, N.; Milekovic, T.; Shkorbatova, P.; Pavlova, N.; Musienko, P.; Bezard, E.; Bloch, J.; Courtine, G. Configuration of electrical spinal cord stimulation through real-time processing of gait kinematics. Nature Protocols 2018, 13, (9), 2031–2061.

40. Merrill, D. R.; Bikson, M.; Jefferys, J. G. R. Electrical stimulation of excitable tissue: design of efficacious and safe protocols. Journal of Neuroscience Methods 2005, 141, (2), 171–198.

41. Rattay, F.; Minassian, K.; Dimitrijevic, M. R. Epidural electrical stimulation of posterior structures of the human lumbosacral cord: 2. quantitative analysis by computer modeling. Spinal Cord 2000, 38, (8), 473–489.

42. Wenger, N.; Moraud, E. M.; Gandar, J.; Musienko, P.; Capogrosso, M.; Baud, L.; Le Goff, C. G.; Barraud, Q.; Pavlova, N.; Dominici, N.; Minev, I. R.; Asboth, L.; Hirsch, A.; Duis, S.; Kreider, J.; Mortera, A.; Haverbeck, O.; Kraus, S.; Schmitz, F.; DiGiovanna, J.; van den Brand, R.; Bloch, J.; Detemple, P.; Lacour, S. P.; Bézard, E.; Micera, S.; Courtine, G. Spatiotemporal neuromodulation therapies engaging muscle synergies improve motor control after spinal cord injury. Nature Medicine 2016, 22, 138.

43. Bickel, C. S.; Gregory, C. M.; Dean, J. C. Motor unit recruitment during neuromuscular electrical stimulation: a critical appraisal. European Journal of Applied Physiology 2011, 111, (10), 2399.

44. Lucas, E.; Whyte, T.; Liu, J.; Russell, C.; Tetzlaff, W.; Cripton, P. A. High-Speed Fluoroscopy to Measure Dynamic Spinal Cord Deformation in an In Vivo Rat Model. J. Neurotrauma 2018, 35, (21), 2572–2580.

45. Kathe, C.; Skinnider, M. A.; Hutson, T. H.; Regazzi, N.; Gautier, M.; Demesmaeker, R.; Komi, S.; Ceto, S.; James, N. D.; Cho, N.; Baud, L.; Galan, K.; Matson, K. J. E.; Rowald, A.; Kim, K.; Wang, R.; Minassian, K.; Prior, J. O.; Asboth, L.; Barraud, Q.; Lacour, S. P.; Levine, A. J.; Wagner, F.; Bloch, J.; Squair, J. W.; Courtine, G. The neurons that restore walking after paralysis. Nature 2022, 611, (7936), 540–547.

46. Capogrosso, M.; Wenger, N.; Raspopovic, S.; Musienko, P.; Beauparlant, J.; Luciani, L. B.; Courtine, G.; Micera, S. A computational model for epidural electrical stimulation of spinal sensorimotor circuits. Journal of Neuroscience 2013, 33, (49), 19326–19340.

