## Supplementary Information for "Injectable ventral spinal stimulator evokes programmable and biomimetic hindlimb motion"

**This file includes:**

Materials and Methods

Figs. S1 to S16

Captions for movies S1 and S5

References

**Other supplementary material:**

Movies S1 to S5

**Materials and Methods**

**VentrE design**

VentrE consists of 3 regions: spinal interface, stem, and I/O (**Fig. S1A**). The spinal interface region has a total length, L_si_ = 14 mm, 0.1 mm diameter stimulation electrodes were deployed at 9 evenly distributed positions along the longitudinal axis. The stem region has a total length, L_stem_ = 68 mm, that is *ca.* twice the length between the T10 vertebra and skull of an adult mouse ^1^ with ca. 35 mm to allow full motion in freely behaving mice without stretching the stem or breaking electrical connections. Both longitudinal and transverse elements have a wave-like structure consisting of alternating semicircles with periodicity, L_w_ = 0.1 mm (semicircle diameter = 0.05 mm) with all elements having widths, W_ribbon_ = 10 μm. This structure helps to accommodate strains associated with spinal cord bending in normal freely behaving animals as reported previously ^2^. The pitches of the longitudinal and transverse elements, P_long_ and P_trans_, are 0.1 and 1 mm. respectively and the total width of the spinal interface, W_si_, is 0.85 mm. We included six redundant independently-addressable stimulation electrodes at positions 3 – 8 where the distance between redundant electrodes is L_bk_ = 0.15 mm is ca. 10x shorter than the ca. 1.6 mm distance between the nine stimulation sites. The redundant electrode pairs have independent parallel Au interconnects, 1.5 μm in width routed along their respective longitudinal elements (**Fig. S1C**) as described previously ^3^ with 15 I/O pads allowing each electrode to be independently addressed (**Fig. S1A**, The stimulation electrodes are double-sided platinum (Pt) stimulation electrodes with a diameter, D_e_ = 0.1 mm, and the I/O pads are L_pad_ = 0.2 mm in length and W_pad_ = 0.1 mm in width, with rhomboidal open windows that allow reliable electrical bonding as described previously ^4^.

**Fabrication of VentrE**

The VentrE fabrication is similar to our previous reports for implantable neural probes ^4, 5^{Lee, 2019 #14}{Lee, 2019 #32}. The fabrication of VentrE includes 5 layers of photolithography: (i) Deposition of the bottom Pt electrode layer; (ii) Spin-coating and photolithography (PL) definition of the bottom polymeric backbone/encapsulation layer; (iii) Deposition of the Au interconnect layer; (iv) Spin-coating and PL definition of the top polymeric backbone/encapsulation layer; (v) Deposition of the top Pt electrode layer. Details of the step-by-step fabrication procedures can be found in Fig. S2B. The fabrication of the I/O interface in the same device involves two additional steps: (i) Deposition of a bottom Au layer; (ii) Spin-coating and PL definition of a thin polymeric supporting layer for I/O pads. Details can be found in Fig. S2C. For the deposition of metal layers, Shipley 1805 (Microposit, The Dow Chemical Company, USA) positive photoresist was used to define the pattern prior to deposition; while the SU-8 2000.5 photoresist (MicroChem Corp., USA) was spin-coated and defined by PL for the polymeric backbone/encapsulation. The fabricated VentrEs are released from the Si wafer fabrication substrates by etching a Ni sacrificial layer with in-house made etchant (1:1:20 of 40% FeCl_3_ (Sigma Aldrich, USA), 37% HCl (Sigma Aldrich, USA), and DI water) for 3-5 hrs, rinsed in DI water 5 times, and transferred to sterile 1× phosphate-buffered saline (PBS) solution (HyClone™ Phosphate Buffered Saline, Thermo Fisher Scientific, USA) for later use.

**Estimation of the spinal cord surface strain upon bending**

Here, the spinal cord model was considered as a symmetric structure so that the length of its central line remains unchanged upon bending. Although the spinal cord exhibits natural curvatures, we assumed that it is straight in its relaxed state.

Under the above assumptions, the surface tensile strain of the spinal cord can be calculated as:

$$\sigma=\frac{L_{t}-L_{0}}{L_{0}}= \frac{\frac{\pi\left( d+T_{sp} \right)}{2}-\frac{\pi d}{2}}{\frac{\pi d}{2}}=\frac{T_{sp}}{d}\times100\%$$

where *L_0_* is the original length of the spinal segment; *L_t_* is the surface length of the spinal segment after bending; *T_sp_* is the thickness of the spinal cord, and *d* is the diameter of the curvature (**Fig. S3I**). Here, the *T_sp_* was calculated by measuring and averaging the thicknesses at the lumbar enlargements (maximum in the lumbosacral segment) from 5 adult CD-1 female mice, affording a value of *ca.* 1.85 mm.

**Finite element analysis of strain distribution**

COMSOL Multiphysics® was used to simulate the strain distribution in a serpentine VentrE segment under tension and compression. In the physical model, the VentrE segment has a total thickness of 600 nm (SU8), with a 100-nm Au metal layer embedded in the center (**Fig. S4B and C**). The VentrE segment was bonded to the surface of a bulk PDMS substrate (Length × Width × Height = 0.3 mm × 0.1 mm × 0.05 mm), which mimics the dura mater of the spinal cord (**Fig. S4A**). A tensile/compressive displacement of either 15% or 30% was imposed on the PDMS substrate and the maximum strains in the SU8 and the embedded Au layer were calculated accordingly.

**Tensile elongation and fatigue tests.**

For the tensile elongation and fatigue tests, a modified serpentine structure was used (**Fig. S5**), which has the same wave-like periodicity metal interconnects as the VentrE across the entire length, with 100 μm diameter pads incorporated at both ends of the longitudinal elements to enable resistance measurements. For the tensile elongation test, the serpentine device structure was deposited on PDMS, which was then clamped and stretched to various displacements in an optical microscope to image the structure as a function of percentage elongation (**Fig. S6**). For fatigue tests, the devices were affixed to a PDMS substrate clamped at both ends to a custom-designed and motorized uniaxial linear stretching device (**Fig. S7**) that allows repeated stretch/relaxation cycles. The resistance of individual longitudinal Au interconnects was measured using a probe station (Keysight Technologies, USA) directly on the PDMS without further transfer. The resistance was either measured in the stretched state (**Fig. S8I**), or in the relaxed state after fatigue cycles (**Fig. S7, Fig. 1D**). During fatigue tests, a tensile strain of *ca.* 18% was imposed on the PDMS, which transfers a similar nominal strain to the serpentine devices.

**Vertebrate animal subjects**

Adult (24-30 g) female CD-1 mice (Charles River Laboratories, USA) were used in this study as the vertebrate animal subjects. All procedures conducted on the mice were approved by the Animal Care and Use Committee of Johns Hopkins University and of Harvard University. The animal care and use programs at Johns Hopkins University and Harvard University meet the requirements of the Federal Law (89-544 and 91-579) and NIH regulations and are also accredited by the American Association for Accreditation of Laboratory Animal Care (AAALAC). Animals were group-housed on a 12 h:12 h light:dark cycle in either the Johns Hopkins University Homewood Central Facility (Mudd Hall) or the Harvard University's Biology Research Infrastructure (BRI), and fed with food and water *ad libitum* as appropriate.

**Surgery**

Surgeries were performed under standard aseptic procedures, where autoclavable tools were autoclaved for 1h before use and the devices and other plastic tools are disinfected in 70% ethanol and washed in sterile 1x PBS before use. The sterile device was loaded into a polyimide micro-catheter tube (Nordson MEDICAL, USA; ID: 0.008’’; OD: 0.0095’’) using an injector. The loading procedures were detailed in our previous report ^6^. Once VentrE was loaded, the injector was then connected to a syringe pump for precise saline injection. The polyimide (PI) tube was inserted in a second polyethylene tube (SAI Infusion Technologies, Inc; ID: 280 μm; OD: 635 μm) with a length of *ca.* 5.8 cm for protection of the VentrE’s stem (**Fig. S11**). The injection setup was mounted onto a stereotaxic frame (Lab Standard Stereotaxic Instrument, Stoelting Co., USA). CD-1 mice were anesthetized by intraperitoneal injection of a mixture of 75 mg/kg of ketamine (Patterson Veterinary Supply, USA) and 1 mg/kg dexdomitor (Orion Corporation, Finland). The degree of anesthesia was verified by the toe-pinch method prior to the surgery. Anesthetized mice were placed on a 37 °C homeothermic blanket (Harvard Apparatus, USA) to maintain their body temperature.

Before the implantation surgery, a head stage with flexible printed circuits (FPC) electrical interface was first mounted. The shaved and disinfected scalp was incised to expose a *ca.* 6 mm × 8 mm area of the skull. A 1-mm diameter burr hole was drilled on the skull at *ca.* 1.0 mm anteriorly and *ca.* 1.0 mm laterally to Bregma. A sterilized 0-80 set screw (18-8 Stainless Steel Cup Point Set Screw; outer diameter: 0.060″ or 1.52 mm, groove diameter: 0.045″ or 1.14 mm, length: 1/4″ or 6.35 mm; Fastenere, USA) was screwed into the hole to a depth of *ca.* 800 μm as the reference electrode. The 3D printed head stage (**Fig. S4**) with a pre-mounted FPC electrical interface was then fixed to the skull using dental cement (METABOND, Parkell, USA).

The surgical subject was fixed on a custom-designed rotatory stage. A *ca.* 1.5-cm skin incision was made on the animal’s back, from *ca.* the T8 to L2 vertebrae. The T13 vertebrae were identified (i.e., the attached rib as the landmark) and the T9-T13 vertebrae were then exposed by dissecting the muscles from the spinous processes. The exposed vertebrae were cleaned and dried. (ii) the rotatory stage was fixed in the stereotaxic frame and the animal was brought to 60-75° with respect to the horizontal direction. (iii) the PI microcatheter tube was inserted into the spinal canal through a gap between two vertebrae (either the T10/T11 gap for ventrolateral implantation or the T11/T12 gap for dorsal implantation). The depth of insertion is *ca.* 2 cm. (iv) The PI tube was then retracted together with a pulse of saline at 60-80 ml/hr with a duration of 2-5 seconds. The saline flow was stopped once the VentrE in the tube was moved steadily. The tube was continuously retracted until it is fully out of the spinal canal. (v) The VentrE was gently pulled out until the landmark on the stem (**Fig. S2A**) appeared, indicating the correct rostrocaudal positioning of the VentrE. A representative implantation procedure is shown in **Movie S2**.

Following implantation, the interfaces were secured using the procedures shown in **Fig. S11.** Briefly, the PE protection tube was fixed to the vertebrae near the entrance of the VentrE by dental cement; the I/O pads were aligned onto the FPC on the head stage as reported previously ^4^; and **t**he other end of the PE tube was fixed to the skull of the subject using dental cement. After the I/O interface dried, epoxy was applied on top for encapsulation and the remaining portion of the probe was further secured with dental cement.

Completion of surgery. Once the interface was secured, incisions were closed using tissue adhesive (Vetbond^TM^, 3M, USA), and antibiotic ointment (Acme United Corporation, USA) was applied to prevent infection.

*Sham surgery*

For the sham surgeries, all the procedures were performed on mice as with the VentrE implantation surgery described above, except for the actual implantation of VentrE.

**Surgery recovery evaluation**

An open-field mobility test was employed to evaluate the recovery of mice post-surgery ^7, 8^. A 40 cm × 42 cm platform (**Fig. S12A**) with a height set to *ca.* 25 cm and a black surface was used for video tracking of the white CD-1 mice. The platform was cleaned by 70% ethanol (in DI) and fully dried prior to each trial. The camera (model, manufacturer) was installed *ca.* 80 cm above the platform, and videos were recorded at 30 frames per second (fps) on a computer. The activity of mice was tracked on days 1, 2, 3, 5, and 7 after surgery.

The recorded mice trajectories (**Fig. S12B**) were analyzed offline using a custom Python script that was used to calculate real-time speed (v_real_), maximum speed (v_max_), average speed (v_avg_), distance traveled (d), and the idle time (t_idle_). Except for the idle time, which was refreshed every frame, the other parameters were calculated and refreshed every 10 frames (*ca.* 0.33 s). A mouse was considered idle if its real-time speed was lower than 4 cm/s, which was empirically determined based on control mice without surgery.

**Kinematics**

Colored markers were put on the skin overlying the key joints (iliac crest, hip, ankle, and metatarsophalangeal (MTP) joints) to record hindlimb motion ^9^. The knee joint position was determined by triangulation from the hip and the ankle as described previously to avoid analysis errors due to skin slippage ^9^. The hindlimb motion was captured at 240 fps and analyzed offline by the in-house Python scripts.

**Horizontal ladder walking test**

A horizontal ladder (Maze Engineers, USA) customized for mice (**Fig. S13A**) with a total length of 60 cm, rung pitch of 2 cm, and rung diameter of 3mm (**Fig. S13B**), was used ^10^. Prior to surgeries, mice were trained on the horizontal ladder for 3 days. 15-20 trials were performed per day for each mouse, food rewards were provided at the end of each trial ^11^. Following surgery, the awaked mice were put onto the horizontal ladder and motivated to walk by food rewards. Their walking behavior was recorded at 240 fps to allow detailed offline kinematic analysis. The hindlimb kinematic analyses were performed using an in-house python script.

**Histology**

**Tissue fixation.** Mice were deeply anesthetized and perfused at the end of specific studies using 40 ml 1x PBS at 2 ml/min followed by 40 ml 4% paraformaldehyde in 1x PBS at the same rate. After perfusion, the lumbosacral segments were removed by laminectomy ^12^ and immersed in 4% paraformaldehyde (PFA, in 1x PBS) for 24 hrs to further fix the spinal tissue. The tissue was then washed in 1x PBS for 24 hrs to remove unreacted PFA. The rostral and caudal lumbosacral segments were sliced using a microtome (Leica, Germany) along the coronal plane at a thickness of 60 μm. The as-obtained coronal sections were incubated in 2% Triton x-100 (in 1x PBS) for 24 hrs to increase the membrane permeability.

**Immunohistochemistry.** Microglia, astrocytes, and neurons were visualized by immunohistological staining against the glial fibrillary acidic protein (GFAP), ionized calcium-binding adapter molecule 1 (Iba1), and neuronal nuclei (NeuN), respectively. The representative lumbosacral coronal sections were incubated in serum-containing anti-GFAP (1:300, Abcam, USA), anti-Iba1 (1:300, Abcam, USA), and anti-NeuN (1:300, Abcam, USA) for 48 hrs. Secondary antibodies labeled with Alexa fluor ® 405, 488, and 647 were used to visualize the immunoreactions with GFAP, Iba1, and NeuN, respectively. The sections were then mounted onto microscope slides and covered with cover glass. A confocal fluorescence microscope (Carl Zeiss Microscopy GmbH, Germany) was used for imaging.

**Quantification of immunostaining.** ZEN (Carl Zeiss Microscopy GmbH, Germany) and arivis Vision4D (arivis AG, Germany) software were used for data visualization. A MATLAB (MathWorks, Natick, MA) code was written for quantitative analysis of immunostaining intensity, which is directly correlated to the distribution of cells ^13^. For unbiased comparison among groups, the intensities of the images were first normalized linearly to the same lowest and highest values. Heat maps of immunostaining intensity were then plotted based on the representative fluorescence images. In the calculation of the heat maps, the original images were divided into regions of interest (ROI) with a size of 10 x 10 pixels. The intensity value of each pixel in an ROI was averaged to yield a single intensity value for the ROI. The calculation was performed for the whole image. The intensity of each ROI in a heat map was normalized to 0-1. The average staining intensity was defined as the average intensity of the ROIs with intensity above 0.03 (0.03 is an empirically determined value below which we believe there is no cell presented in the ROI). All the analyses were conducted blindly.

**Electrical stimulation of the spinal cord**

Electrical stimulation of mouse’s spinal cord was performed in a suspended condition under anesthesia (**Fig. 3a**). In brief, a mouse was suspended and fixed on a customized stage. An Intan^®^ Stimulation/Recording system (Intan, USA) was employed for applying electrical stimuli. The interface mounted on the skull was connected to the amplifier chip of the Intan^®^ system, which was further connected to the controller and computer. Cathode-leading, biphasic current pulses (200 μs per phase) were employed for the study. To obtain the motion amplitude versus current plots (**Fig. S17**), single-pulse stimuli were applied to an electrode at 1s inter-pulse intervals. The current was varied from 1 μA to 700 μA based on the approximate threshold of the site. Train stimulation was employed for the study of triggered motion as well as fatigue tests. Trains of 4-20 pulses at a frequency of 40 Hz were delivered with a 1s inter-train interval. The current was varied from 5 μA to 400 μA to elicit reliable responses from hindlimb muscles. For programmed stimulation, an in-house Python script was used to control the Intan^®^ system. Multiple electrodes were activated in specifically designed sequences to trigger complex motions. The applied current and phase duration was tuned based on the characteristics of each electrode to optimize the resultant motion. Bilateral stimulation was elicited by alternating the stimuli applied to each implanted VentrE stimulator, where the total time per cycle was 0.8 seconds with 0.4 seconds per side. Within the 0.4-second stimulation period, the flexor hotspot was stimulated for 0.1 s, and the extensor hotspot was stimulated for 0.3 s.

**Statistics**

All the statistical evaluations in this work were based on one-way analysis of variance (ANOVA). We assessed *post hoc* pairwise differences by Tukey’s test.

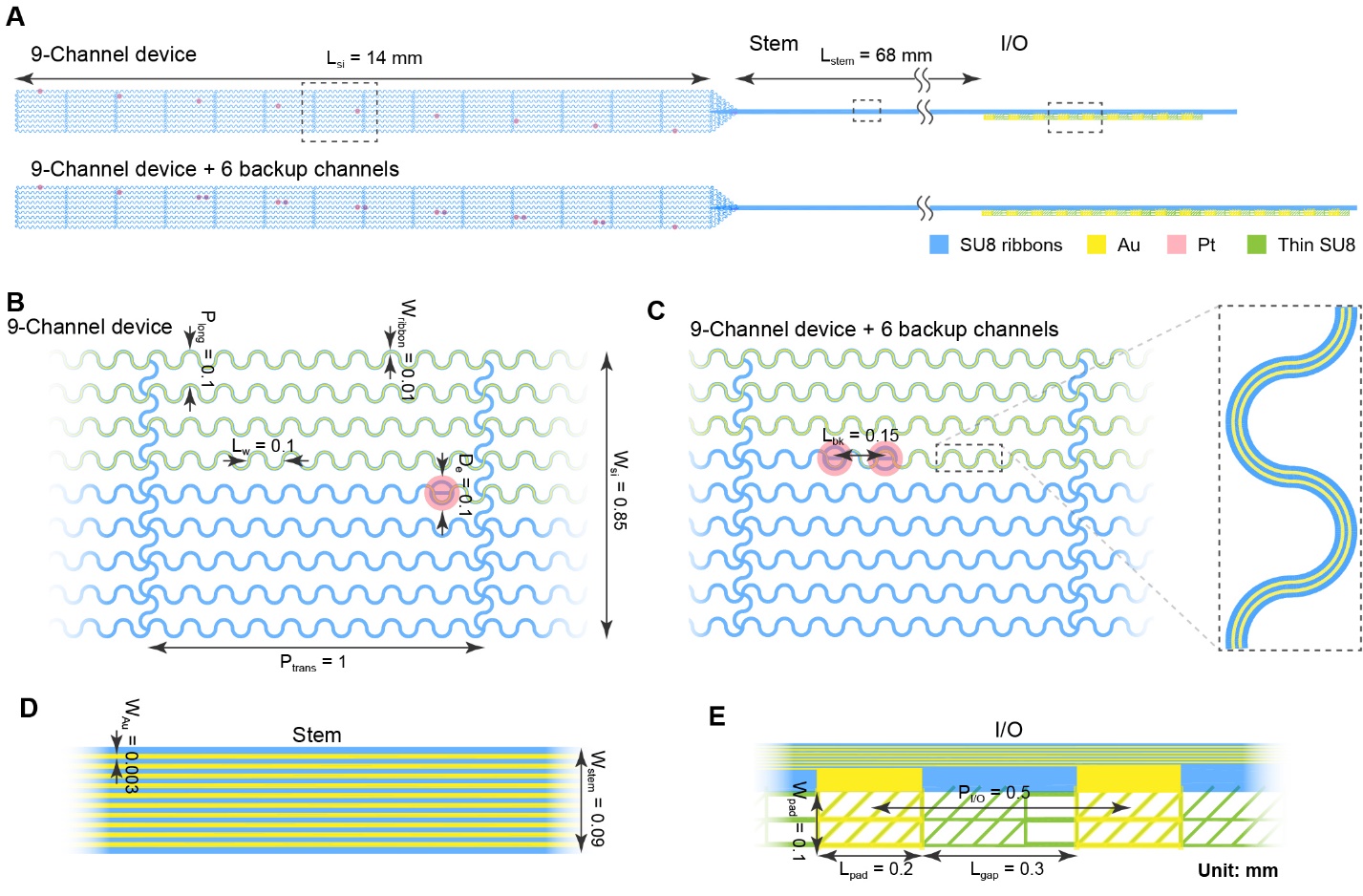

**Fig. S1.** **The VentrE design used in this study.** (**A**) Schematic of VentrE, which consist of 3 regions: the electrode region contains stimulation electrodes; The stem region bridges the electrodes and I/O with passivated Au interconnects; And the I/O region can be further connected to VentrE via a customized interface. VentrE has electrodes positioned at 9 evenly spaced spots along the 14 mm length. For electrical stimulation tests, 6 independently addressable backup electrodes were also included (bottom). (**B**) Magnified illustration of the electrode region of the 9-channel VentrE. The longitudinal and transverse elements have a serpentine structure to accommodate surface strains ^2^. (**C**) Magnified illustration of the electrode region of Design B. The magnified image (*inset*) shows that the two electrodes (one is backup) share the same SU8 ribbon, with two independent Au interconnects embedded in on ribbon. (**D**) Magnified illustration of the stem region. (**E**) Magnified illustration of the I/O. The Au I/O pads (yellow) are supported and connected by a thin layer of SU8 (green).

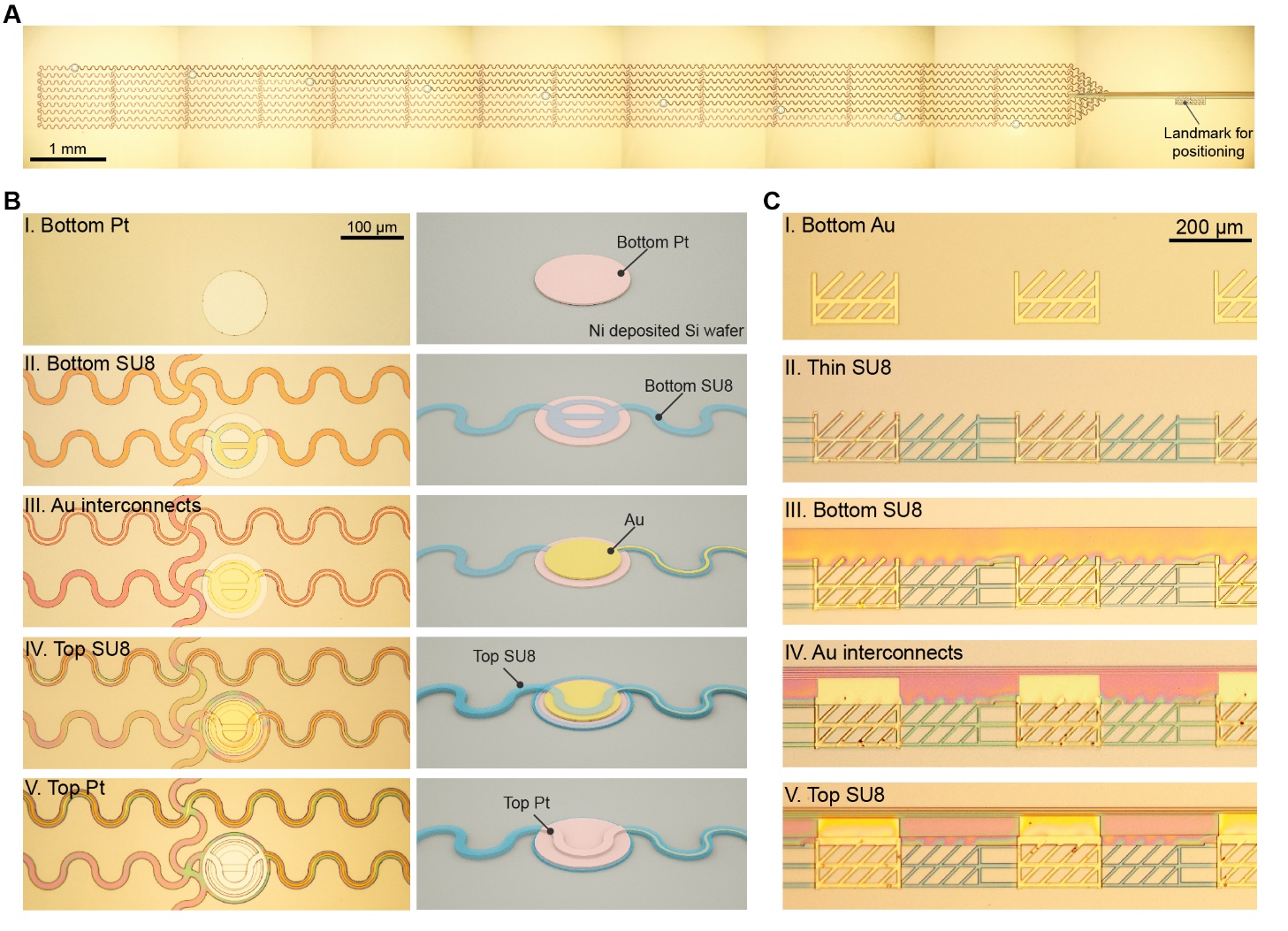

**Fig. S2. Layer-by-layer fabrication of VentrE.** (**A**) Stitched optical microscope image showing the electrode array of a fabricated VentrE. The image was stitched from 8 of 2560×1920 (pixel) images. (**B**) Magnified optical microscope images *(left)* and schematic (*right*) of layer-by-layer fabricated structures on a Ni-deposited silicon wafer in the electrode region. (**C**) Magnified optical microscope images of layer-by-layer fabricated structures on a Ni-deposited silicon wafer at the I/O region.

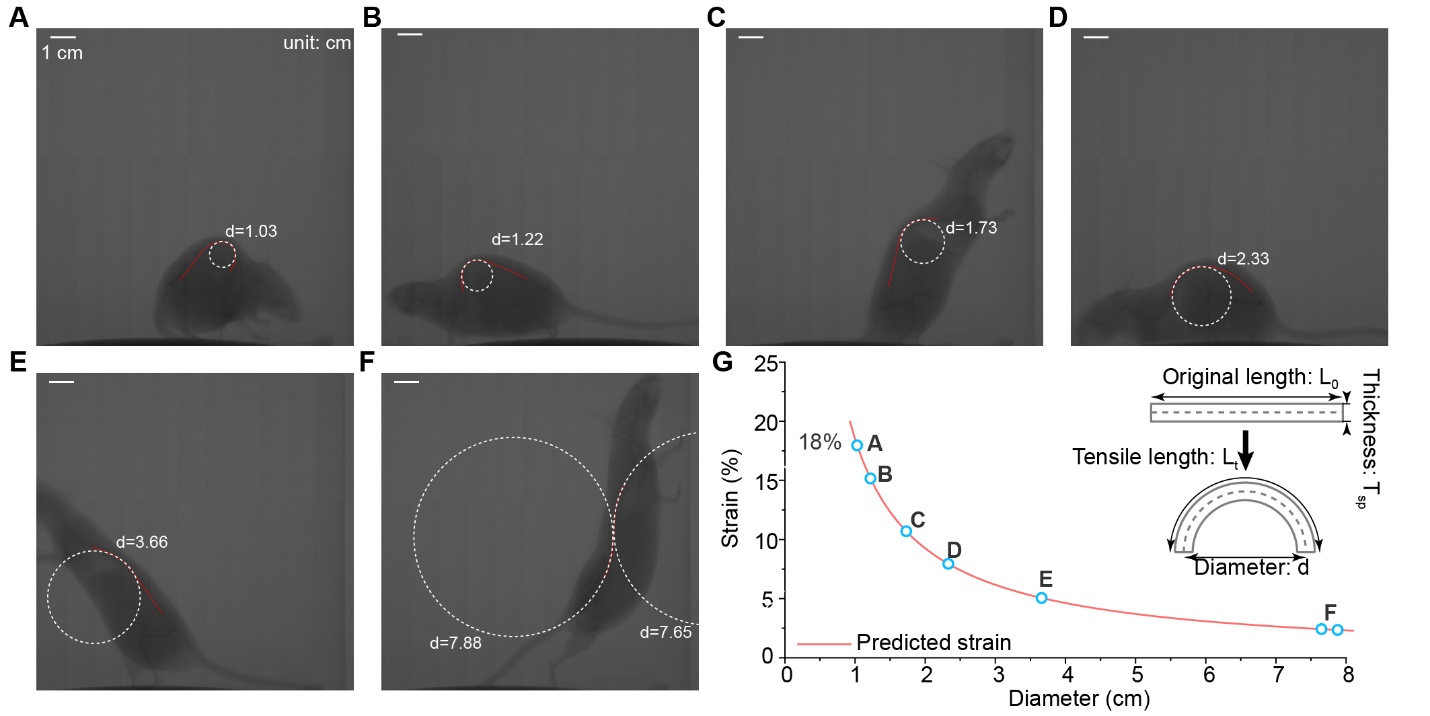

**Fig. S3.** **Estimation of spinal cord surface strain upon bending.** (**A-F**) Representative X-ray images of a freely behaving mouse with various degrees of vertebral column bending. The minimum (**A**) diameters of the vertebral column, which corresponds to the largest strain, were observed to be 1.03 cm. Scale bars: 1 cm; Unit: cm. (**G**) The calculated strain versus diameter curve and the predicted results of various mouse gestures shown in (**A-F**). Inset: a physical model for the prediction of spinal cord surface strain upon bending. The highest predicted strain is *ca.* 18%.

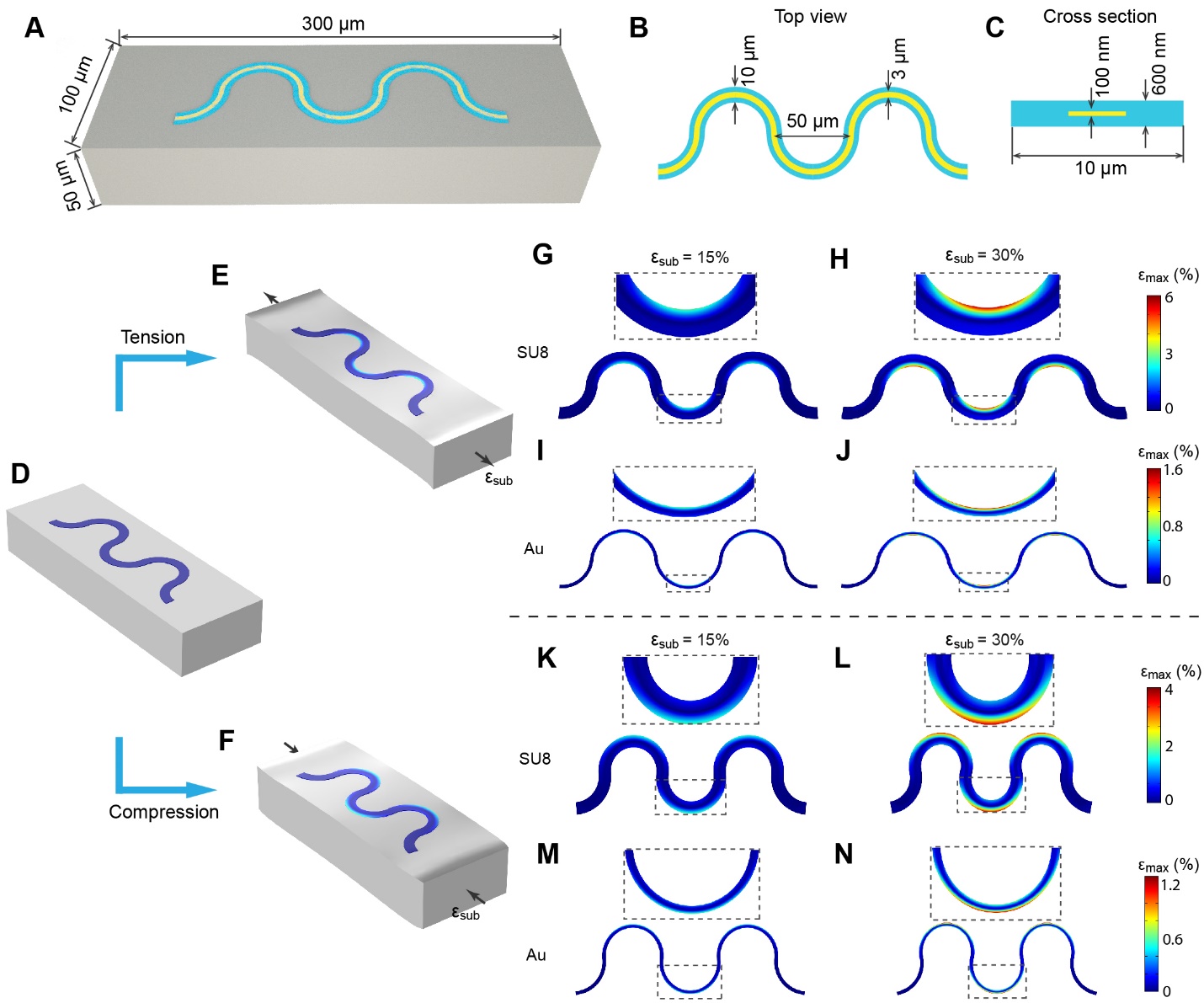

**Fig. S4. Finite element analysis (FEA) of the serpentine device.** (**A**) Physical model employed for FEA: a polydimethylsiloxane (PDMS) substrate with serpentine device segment attached to the top surface is used to mimic VentrE attached to the dura mater of the spinal cord. (**B** and **C**) Top (**B**) and cross-section (**C**) views of the device segment model employed for calculation, with parameters based on the values from the design and fabrication. (**D, E,** and **F**) The model before (**D**) and after (**E** and **F**) strain is applied. Tension (**E**) and compression (**F**) are applied to the substrate to mimic the surface strain produced by the bending of the spinal cord. (**G**-**J**) Distribution of principal strain in SU8 (**G** and **H**) and Au interconnect (**I** and **J**) after 15% (**G** and **J**) and 30% (**H** and **J**) tensile strain are applied to the substrate. (**K**-**N**) Distribution of principal strain in SU8 (**K** and **L**) and Au interconnect (**M** and **N**) after 15% (**K** and **M**) and 30% (**L** and **N**) tensile strain is applied to the substrate.

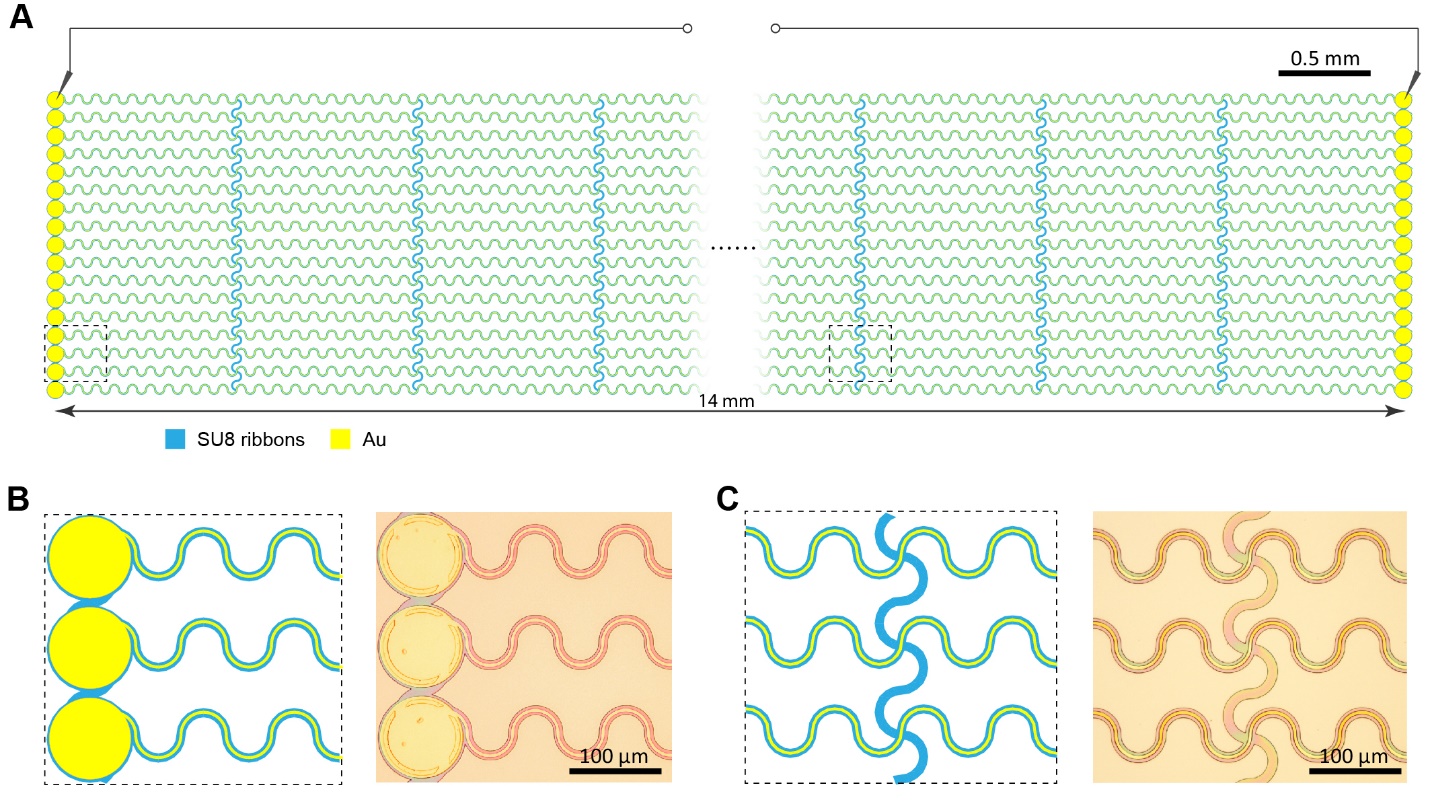

**Fig. S5. Device design for the tensile stretch and fatigue tests** (**A**) Schematic of device used for tensile stretch test. (**B**) The magnified schematic (*left*) and the corresponding optical microscope image (*right*) of the Au electrodes at the end of the device. (**C**) The magnified schematic (*left*) and the corresponding optical microscope image (*right*) of the serpentine region in the middle region of the device.

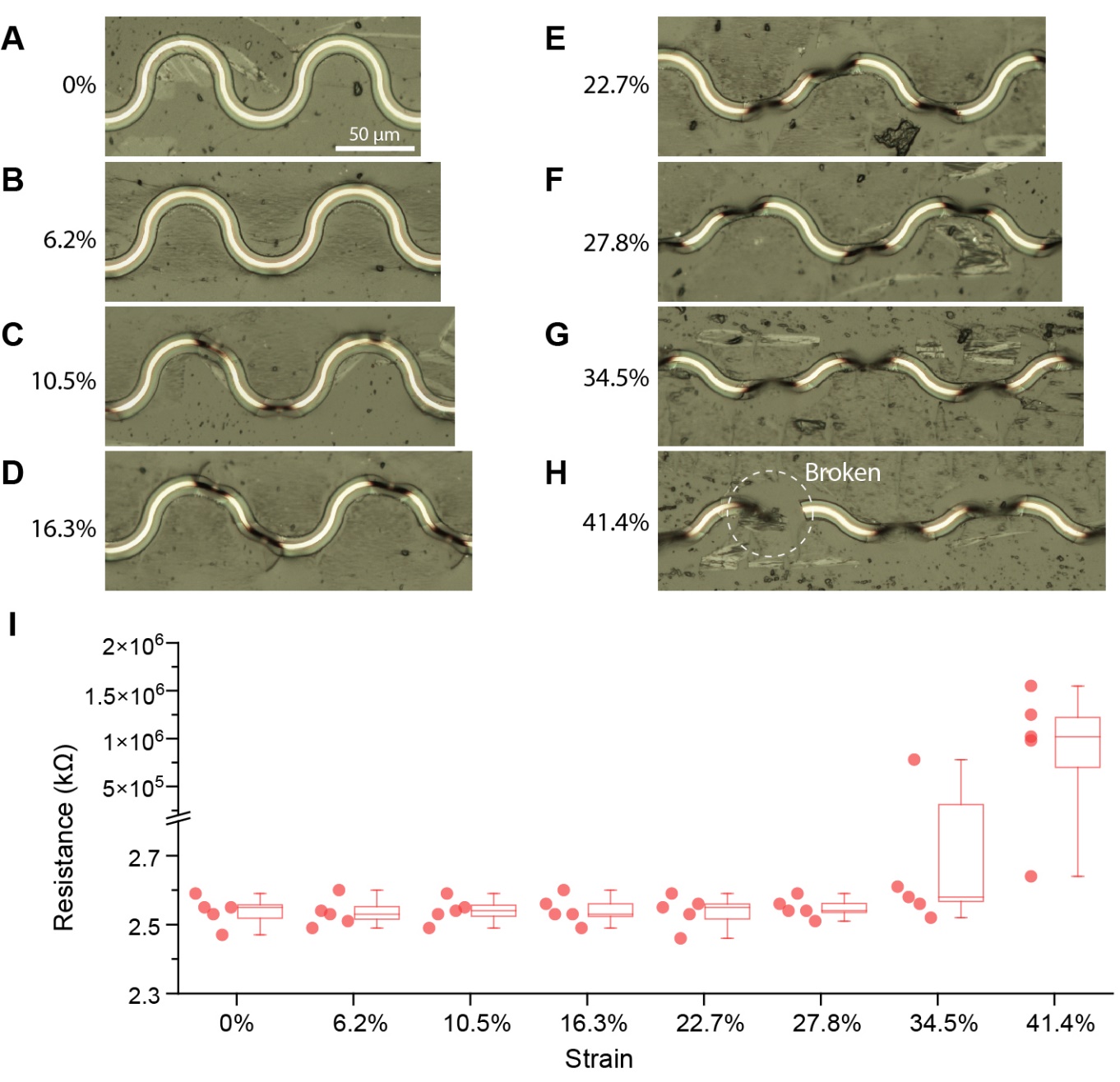

**Fig. S6. Tensile stretch tests of the serpentine ribbons of VentrEs.** (**A**) Optical microscopy image showing the original serpentine VentrE segment without stretching. (**B**-**H**) Optical microscopy images showing the serpentine VentrE segments under tensile strains from 6.2% (**B**) to 41.4% (**H**). The VentrE segment remains continuous to strains as high as *ca.* 34%. (**I**) Box plot showing the resistance at various strains. The measurements were performed when VentrEs were held at the specific strains. The resistance measurements were consistent with the optical observation, where the metal interconnects were mostly connected at a high strain of *ca.* 34%. Error bars: min-max; Box edges: SEM; Lines: median.

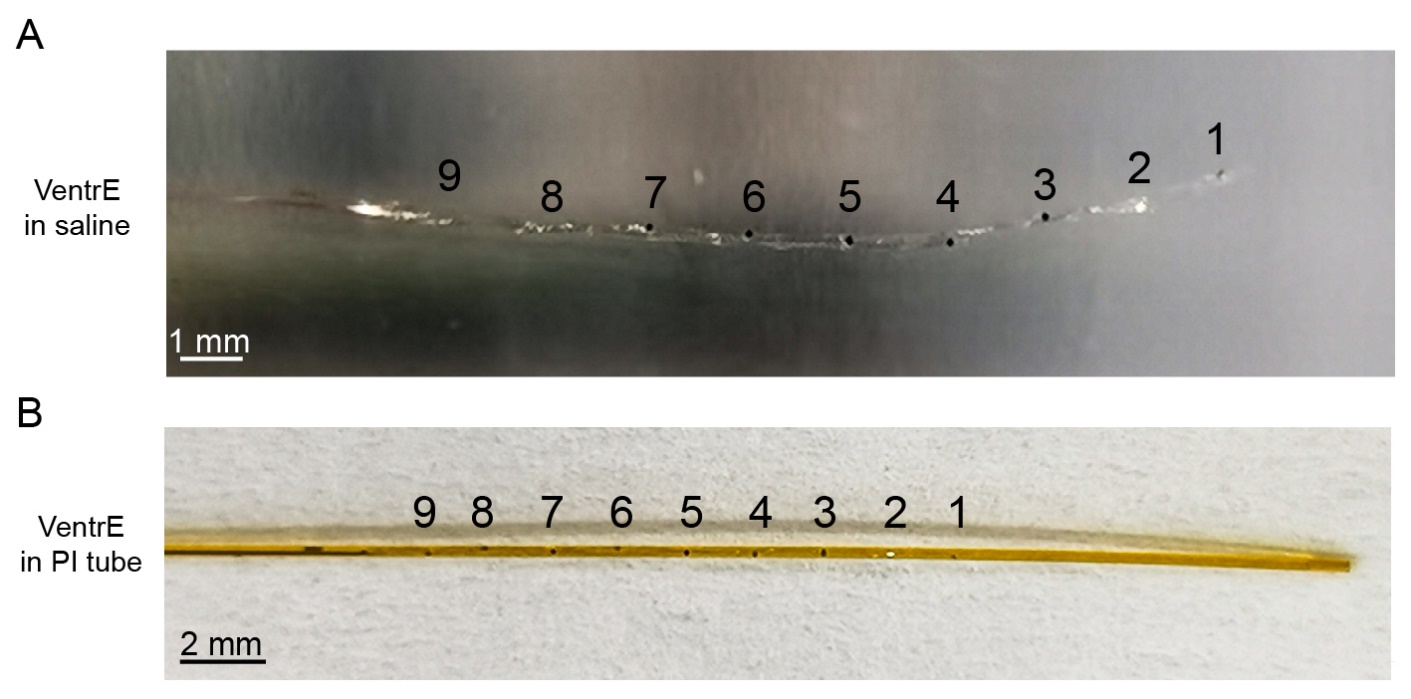

**Fig. S7. The VentrE loading procedure into the polyimide (PI) micro-catheter tube.** (**A**) A VentrE floating in saline was ready to be loaded into a PI tube. (**B**) The PI tube with VentrE loaded. Each electrode can be identified through the transparent PI tube. The indices of the electrodes are labeled accordingly in the images.

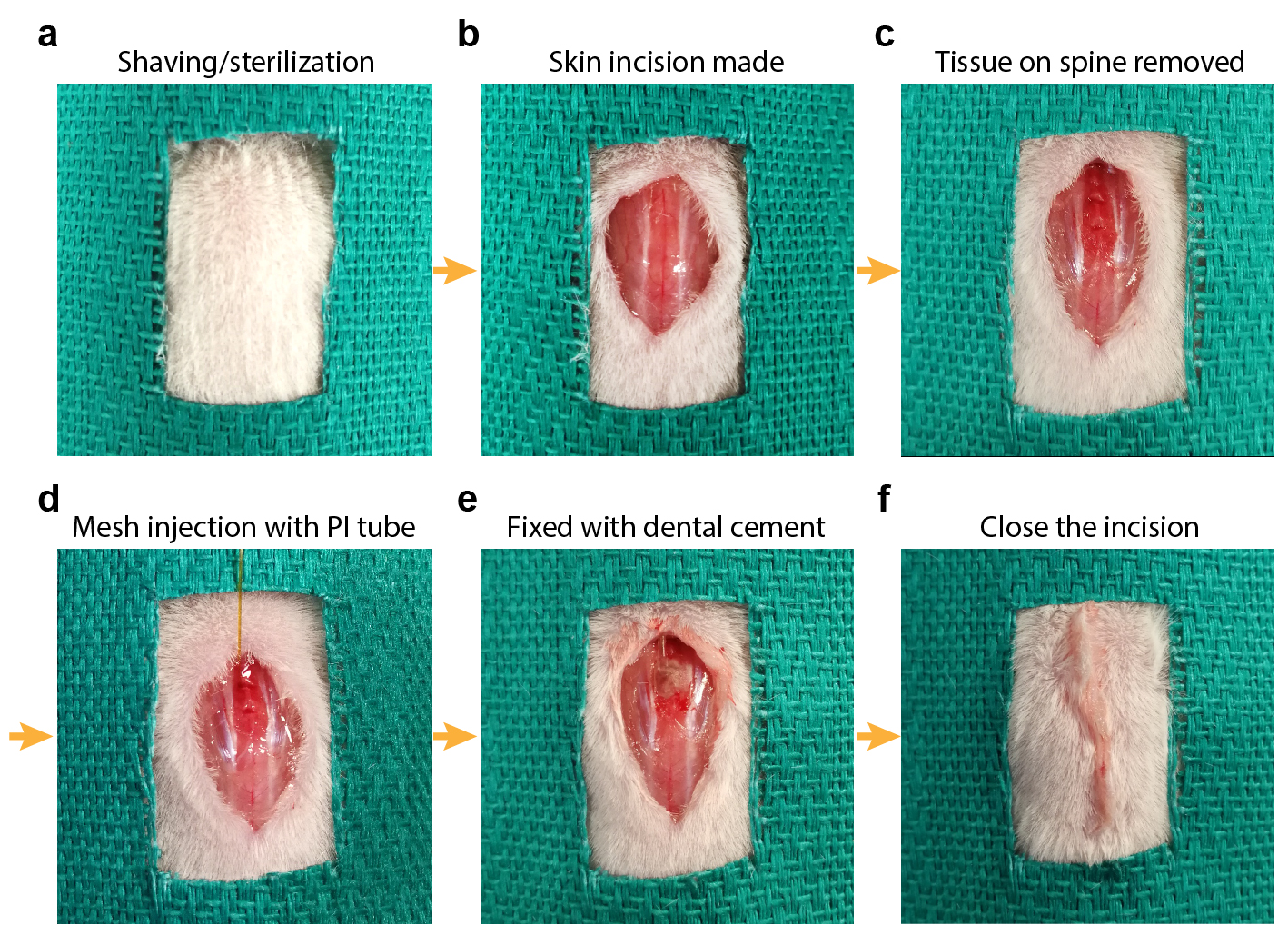

**Fig. S8. The surgical procedures of VentrE implantation.** (**A**) The skin near the implantation region was shaved and sterilized thoroughly. (**B**) An incision was made on the skin near the implantation region approximately from T9 to L2 vertebrae. (**C**) The tissue on the vertebrae is carefully removed to expose the vertebrae and implantation gap. (**D**) A PI microcatheter tube was carefully inserted into the spinal canal through the gap. The insertion point is determined by the target region of VentrE implantation (T10/T11 and T11/T12 gaps for ventrolateral and dorsal implantation, respectively). Once the target depth was reached, the VentrE was ejected from the tube by a saline pulse delivered via a syringe. (**E**) The PI tube was fully extracted, and the PE protection tube was fixed to the spinous processes by dental cement. (**F**) The incision was closed by applying tissue adhesive.

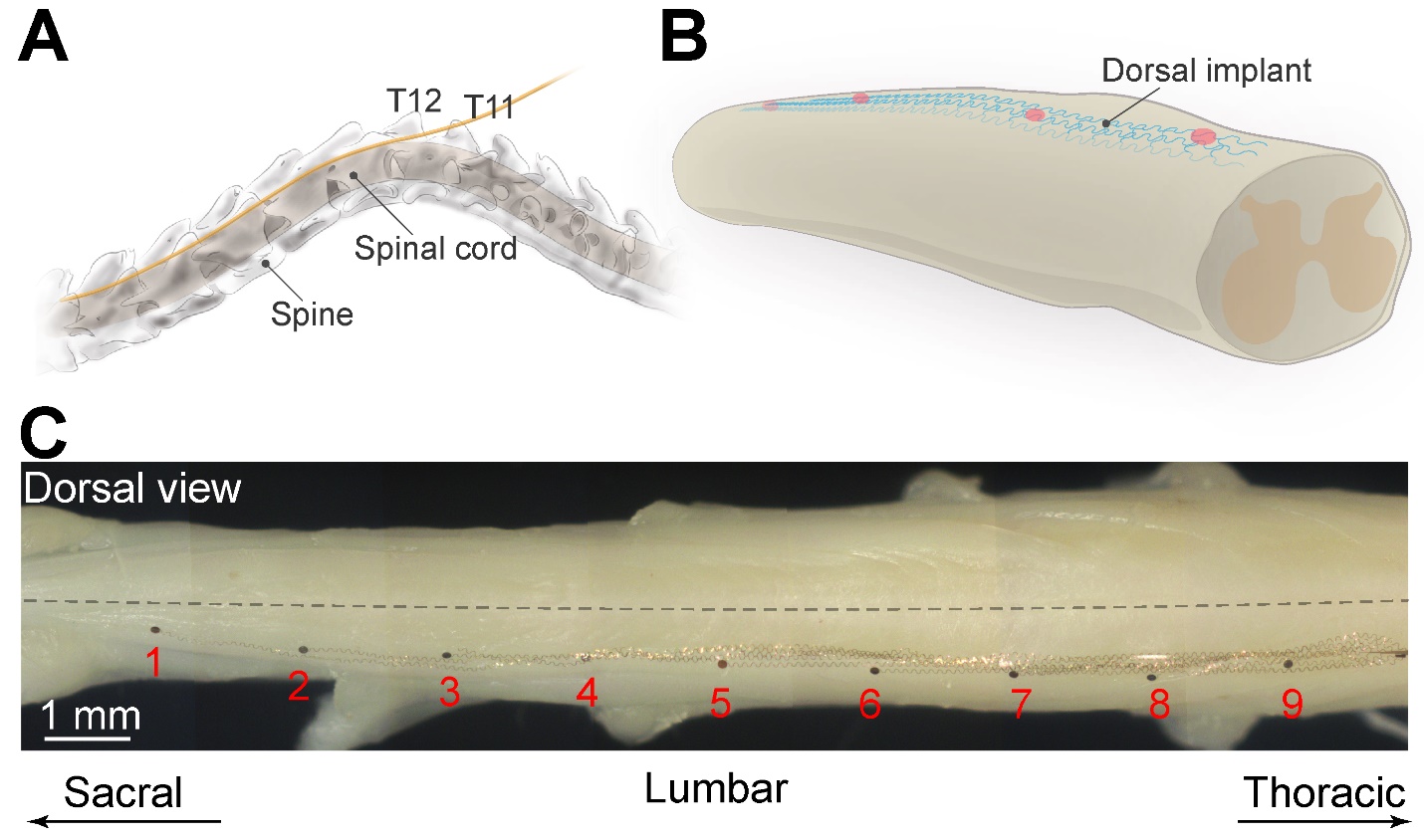

**Fig. S9 Dorsal implantation of the neuroprosthesis. A.** Schematic of the insertion of a microcatheter tube to the dorsal epidural space of the lumbosacral segment. **B.** Schematic illustration of implanted neuroprosthesis on the dorsal epidural surface. **C.** Stitched optical images showing the dorsal epidural surface of the spinal cord with an implanted neuroprosthesis. The electrodes (*1 to 9*) are marked in sequence.

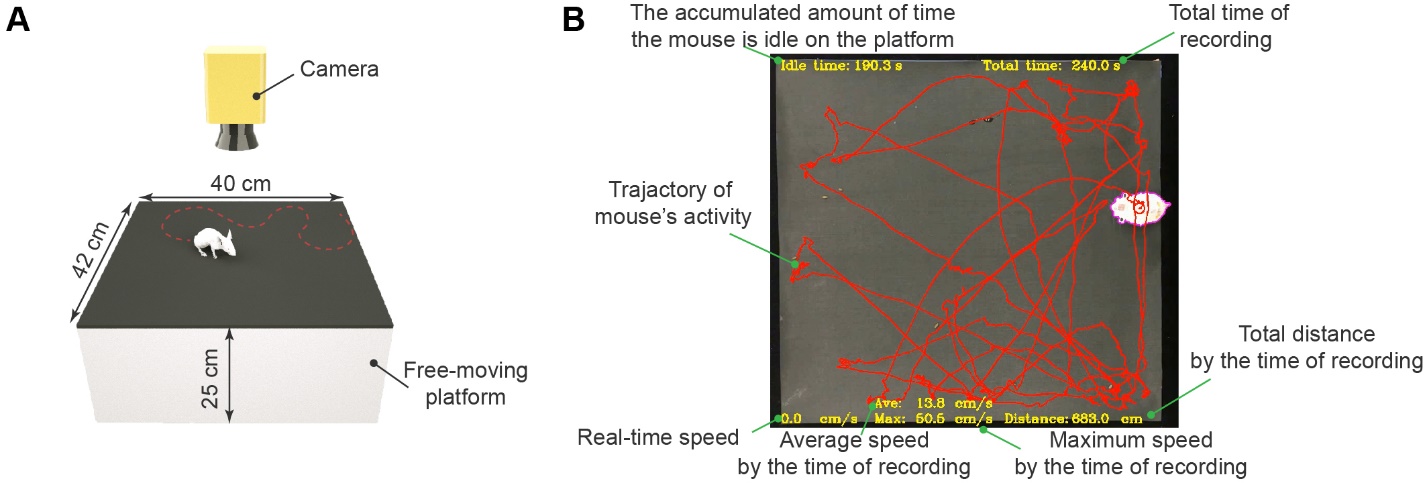

**Fig. S10. Mouse open-field test.** (**A**) Schematic showing the setup for monitoring the unrestrained movement of the mouse before and after surgery. A 42 cm × 40 cm platform with a video camera centered above the platform was used to capture mouse movements for subsequent quantitative analyses (see *Movie S2*). (**B**), A snapshot of a video processed by an in-house Python script for motion tracking. In the algorithm, the idle time, total time, real-time speed, average speed, maximum speed, and total distance are calculated as described in *Supplementary Methods*. The trajectory of the mouse pathway was also recorded and shown (red). Representative videos are included in **Movie S2**.

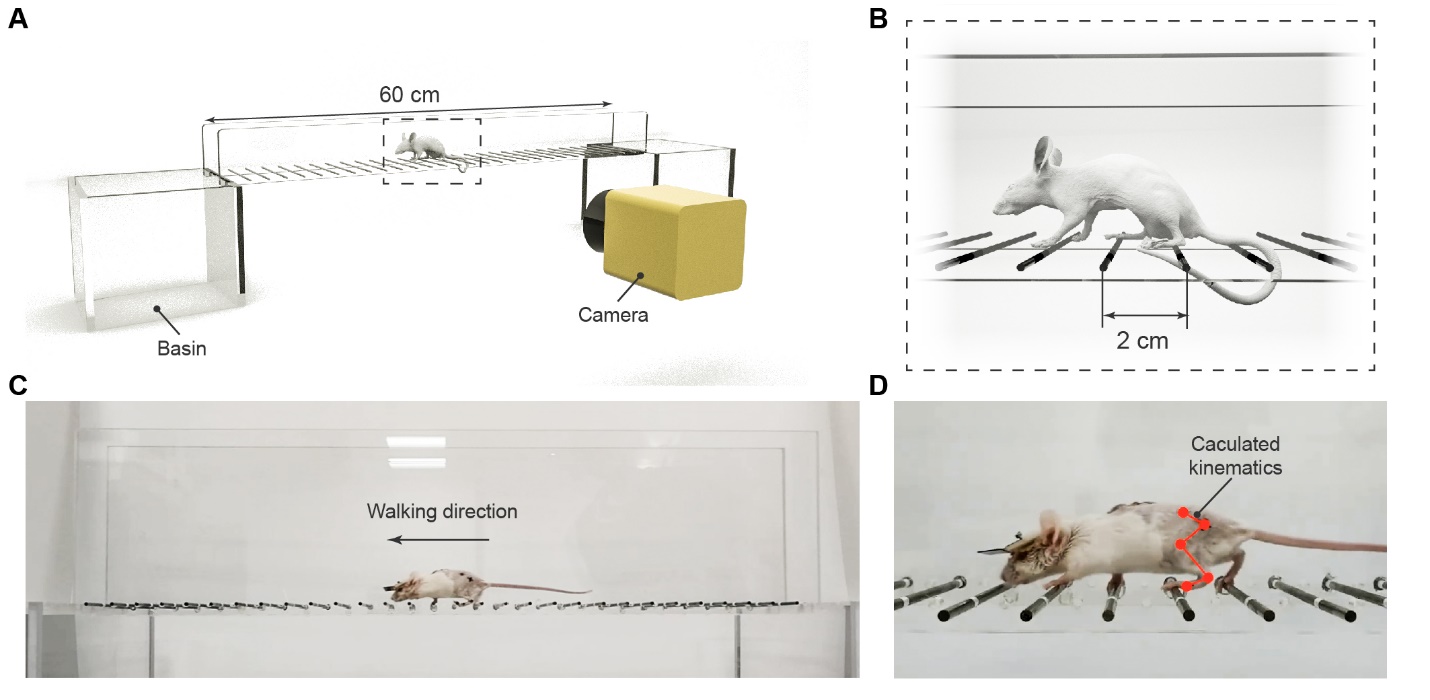

**Fig. S11. The horizontal ladder test.** (**A**) Schematic of the horizontal ladder setup, which contains two basins at both ends of the ladder, a 60 cm total length horizontal ladder, with barriers on either side of the ladder made of transparent acrylic plastic for video recording. (**B**) Magnified schematic of the dashed rectangular region in **A**. The rungs of the ladder have a diameter of 3 mm and a separation between adjacent rungs of 2 cm. (**C**) A snapshot of the video showing a mouse walking on the horizontal ladder. (**D**) A video snapshot showing the calculated kinematics of a mouse walking on the horizontal ladder.

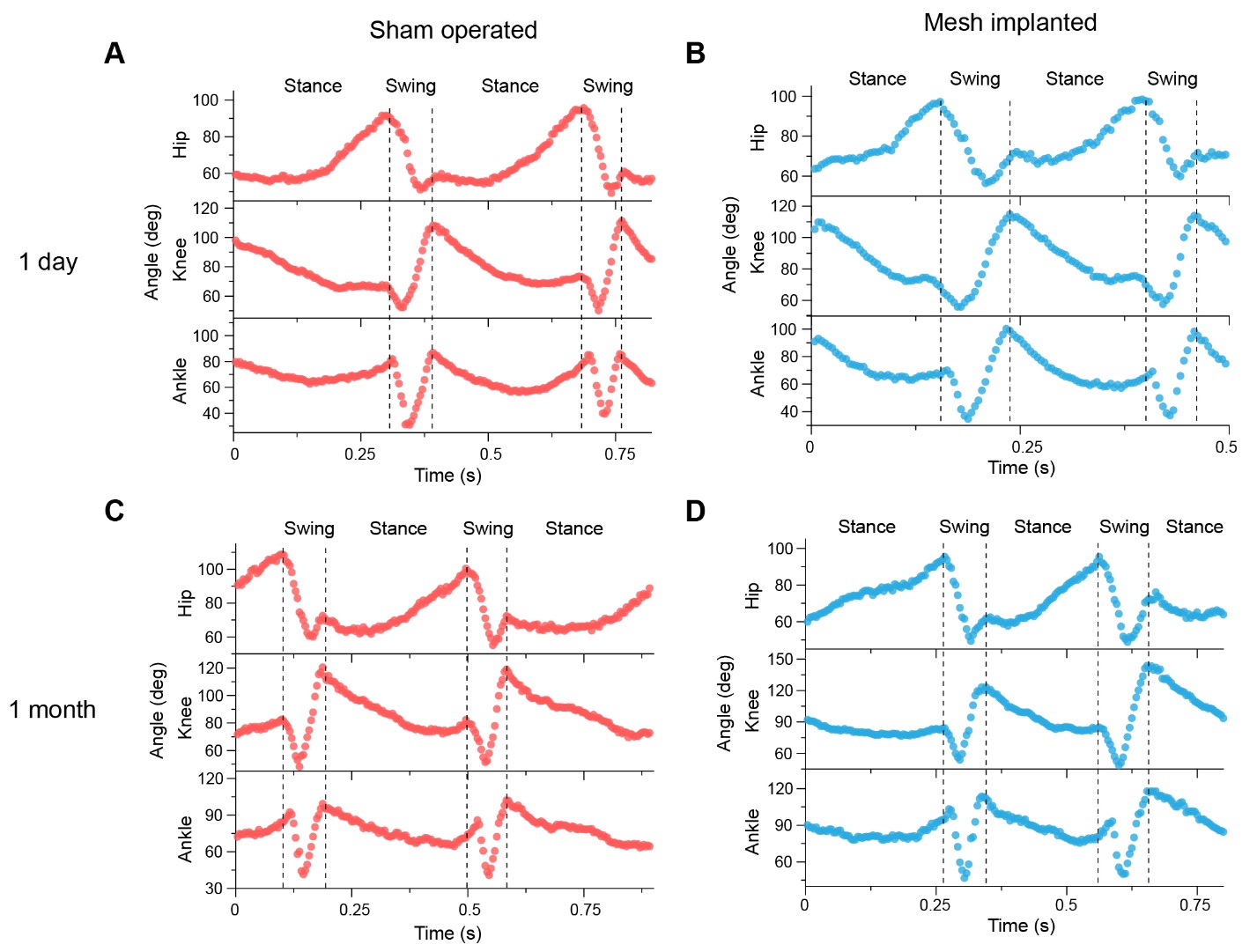

**Fig. S12. Joint angles and phase analysis of mouse hindlimbs when walking on a horizontal ladder.** (**A**) The phase relation of the hip, knee, and ankle joints of a mouse on day 1 after sham-operated surgery. (**B**) The phase relation of the hip, knee, and ankle joints of a mouse on day 1 after VentrE implantation surgery. (**C**) The phase relation of the hip, knee, and ankle joints of a mouse 1 month after sham-operated surgery. (**D**) The phase relation of the hip, knee, and ankle joints of a mouse 1 month after VentrE implantation surgery. 2 step cycles are shown in each plot.

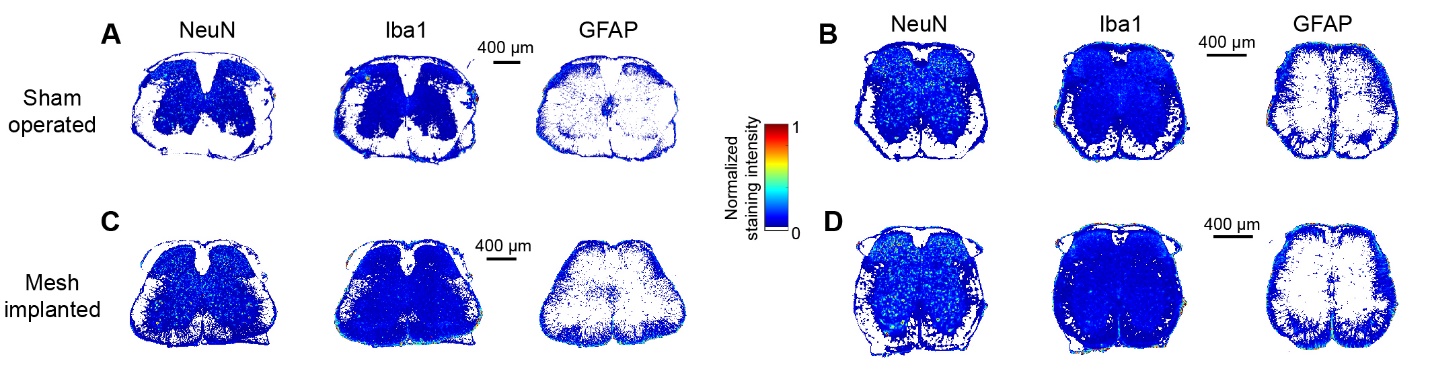

**Fig. S13. Heat maps of the staining intensity of the histology results in Fig. 2.** (**A, B**) Staining intensity maps of the representative lumbar (**A**) and sacral (**B**) coronal sections with sham-operated surgery. (**C, D**) Staining intensity maps of the representative lumbar (**C**) and sacral (**D**) coronal sections with VentrE-implanted surgery. The mice were sacrificed 2 weeks after surgery. The staining intensity maps of NeuN (*left*), Iba1 (*middle*), and GFAP (*right*) are shown accordingly in each panel.

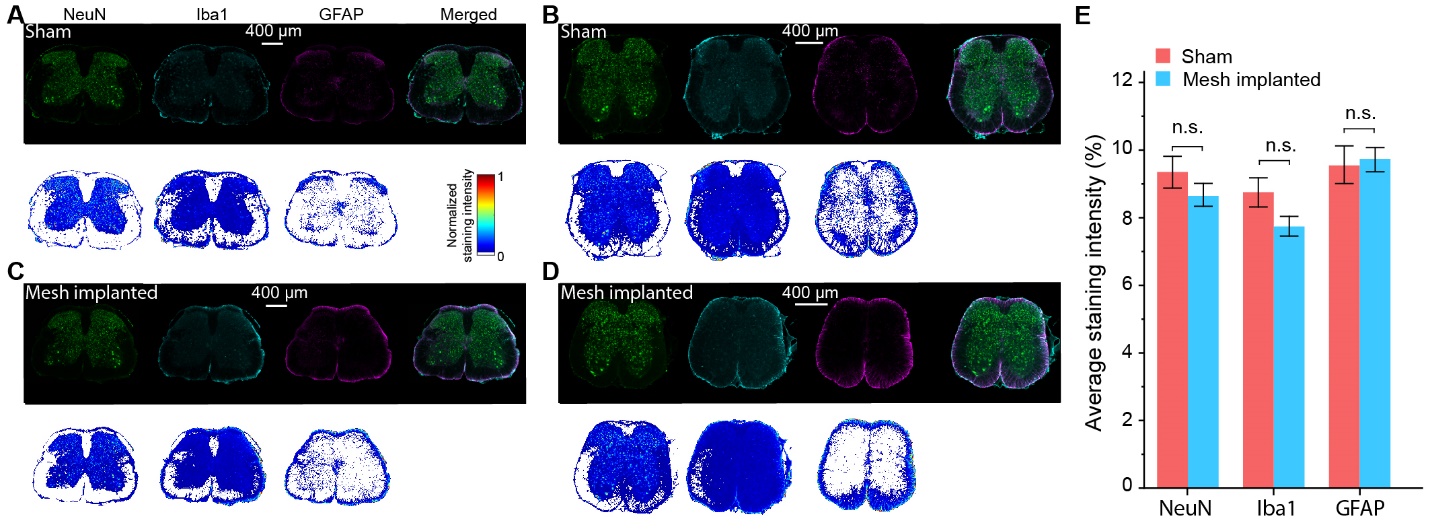

**Fig. S14. Histology studies of the spinal cord lumbosacral segments 4 weeks after surgery.** (**A**-**D**) Fluorescence images (*top*) and the corresponding staining intensity maps (bottom) of the lumbosacral coronal sections with sham-operated (**A** and **B**) and VentrE implantation (**C** and **D**) surgeries. **A** and **C** are from the lumbar segments while **B** and **D** are from the sacral segments. The mice are sacrificed 4 weeks after surgery. Neurons (NeuN, green), microglia (Iba1, cyan), and astrocytes (GFAP, glial fibrillary acidic protein, magenta) are labeled accordingly for each slice. (**E**) Statistic comparisons of the average staining intensities of NeuN, Iba1, and GFAP 4 weeks after surgery (N=3 mice for each condition, 4 slices from each mouse). No significant difference was observed in the three cell types between the sham-operated and VentrE-implanted groups. Error bars: SEM.

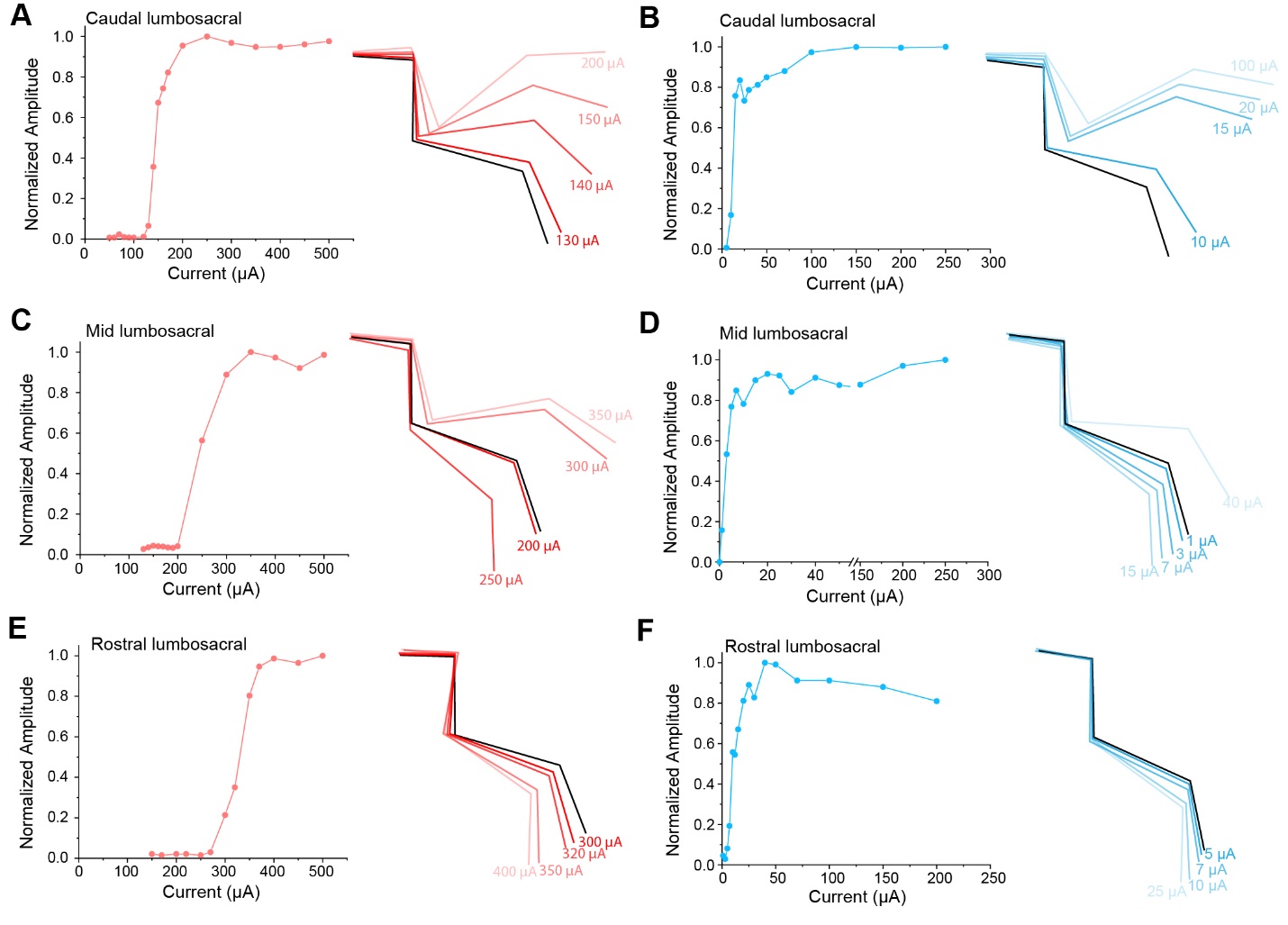

**Fig. S15. Step amplitudes of the hindlimbs versus applied currents at multiple positions of the lumbosacral segment.** (**A** and **B**) Normalized step amplitude versus stimulation current plots (left) and the representative kinematics at various currents (right) when the stimuli were applied to the caudal lumbosacral region. The stimulations on the dorsal (**A**) and ventrolateral (**B**) epidural surfaces are compared. (**C** and **D**) Normalized step amplitude versus stimulation current plots (left) and the representative kinematics at various currents (right) when the stimuli were applied to the mid-lumbosacral region. The stimulations on the dorsal (**C**) and ventrolateral (**D**) epidural surfaces are compared. (**E** and **F**) Normalized step amplitude versus stimulation current plots (left) and the representative kinematics at various currents (right) when the stimuli were applied to the rostral lumbosacral region. The stimulations on the dorsal (**E**) and ventrolateral (**F**) epidural surfaces are compared.

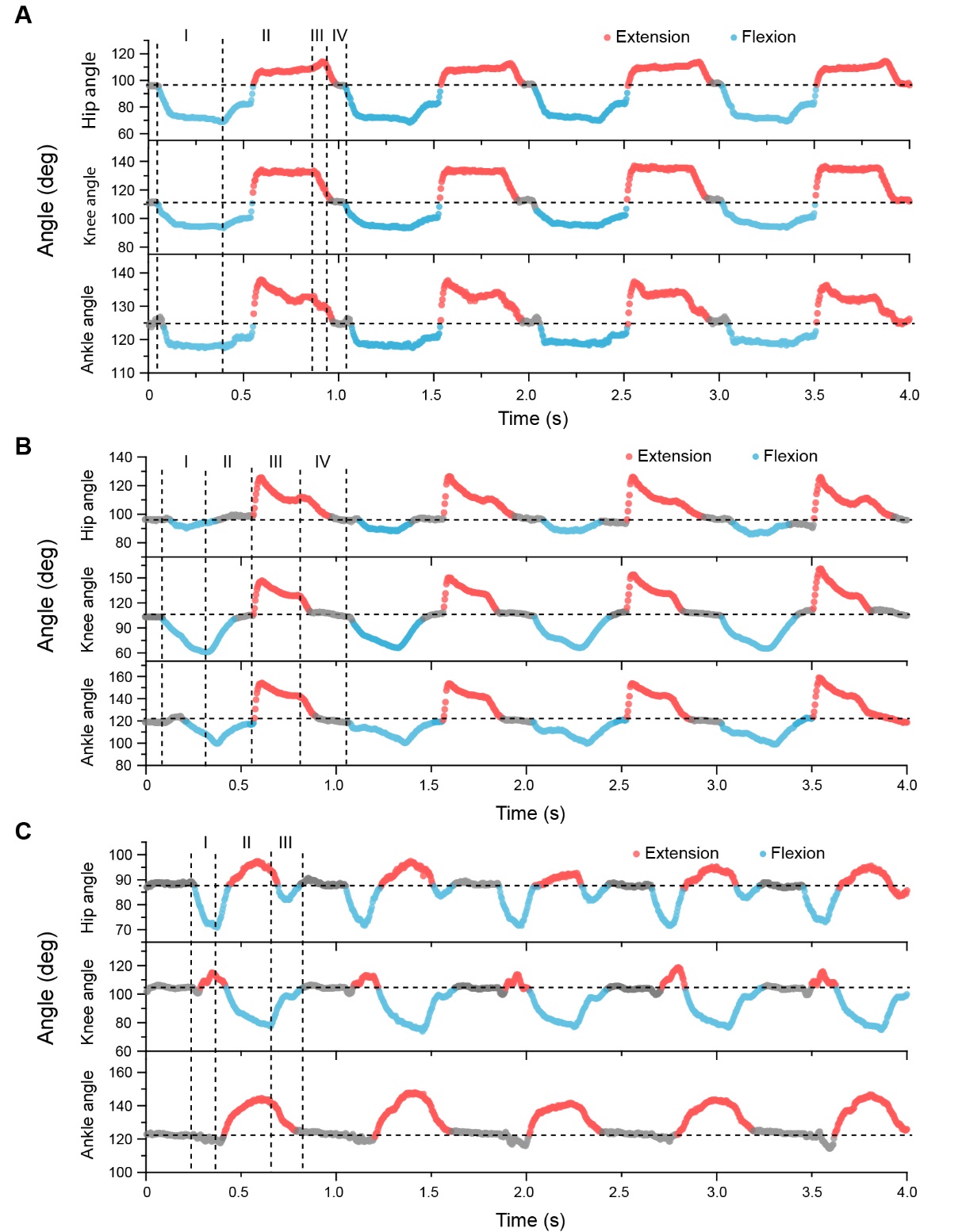

**Fig. S16. Joint angles and phase analysis of mouse hindlimbs under programmed stimulation.** (**A**-**C**) The phase relation of the hip, knee, and ankle joints when the motion was programmed to functionally relevant motion modes of pedaling (**A**), kicking (**B**), and waving (**C**) as presented in **Fig. 4A**, **B,** and **C**, respectively. The extension and flexion of each joint are marked in red and blue dots, respectively, while the grey dots indicate the balanced position.

**Captions of the Supplementary Movies**

**Movie S1. VentrE implantation procedures during surgery.** The representative videos showing the VentrE implantation procedure to both the dorsal and ventrolateral spaces of the mouse spinal canal.

**Movie S2. The activities of mice in the first week after surgery.** The representative videos showing the activities of freely behaving mice on a customized platform since the first day after surgery. The trajectory of their movements, real-time speed, average speed, maximal speed, and total traveling distance are shown instantly on each video.

**Movie S3. The hindlimb kinematics when walking on a horizontal ladder.** The representative videos showing a mouse walking across a horizontal ladder (**Fig. S14**). The representative hindlimb kinematic analyses of mice on day 1 and day 30 after surgery were shown, with either sham-operated or VentrE-implanted surgeries. The videos of kinematic analyses were played at 0.125X speed.

**Movie S4. The programmed stimulation on anesthetized mice.** The videos showed the kinematic analyses of the representative functionally relevant triggered motions (pedaling, kicking, and waving) by programmed stimulation. The videos were played at their original speed.

**Movie S5. The programmed bipedal stimulation.** The videos show the programmed bipedal stimulation enabled by two VentrEs implanted bilaterally to one mouse. The motions and the corresponding kinematics from both the left and right hindlimbs of the mouse were presented. The videos were taken simultaneously from left and right projections. And their timelines were synchronized accordingly for better visualization of the bipedal activities.

**REFERENCES**

1.	Harrison, M.; O'Brien, A.; Adams, L.; Cowin, G.; Ruitenberg, M. J.; Sengul, G.; Watson, C. Vertebral landmarks for the identification of spinal cord segments in the mouse. *NeuroImage* **2013,** 68, 22-29.

2.	Zhang, Y.; Wang, S.; Li, X.; Fan, J. A.; Xu, S.; Song, Y. M.; Choi, K.-J.; Yeo, W.-H.; Lee, W.; Nazaar, S. N.; Lu, B.; Yin, L.; Hwang, K.-C.; Rogers, J. A.; Huang, Y. Experimental and Theoretical Studies of Serpentine Microstructures Bonded To Prestrained Elastomers for Stretchable Electronics. *Advanced Functional Materials* **2014,** 24, (14), 2028-2037.

3.	Fu, T.-M.; Hong, G.; Viveros, R. D.; Zhou, T.; Lieber, C. M. Highly scalable multichannel mesh electronics for stable chronic brain electrophysiology. *Proceedings of the National Academy of Sciences* **2017,** 114, (47), E10046-E10055.

4.	Lee, J. M.; Hong, G.; Lin, D.; Schuhmann, T. G.; Sullivan, A. T.; Viveros, R. D.; Park, H.-G.; Lieber, C. M. Nanoenabled Direct Contact Interfacing of Syringe-Injectable Mesh Electronics. *Nano Letters* **2019,** 19, (8), 5818-5826.

5.	Lee, J. M.; Lin, D.; Hong, G.; Kim, K.-H.; Park, H.-G.; Lieber, C. M. Scalable Three-Dimensional Recording Electrodes for Probing Biological Tissues. *Nano Letters* **2022,** 22, (11), 4552-4559.

6.	Fu, T.-M.; Hong, G.; Zhou, T.; Schuhmann, T. G.; Viveros, R. D.; Lieber, C. M. Stable long-term chronic brain mapping at the single-neuron level. *Nature Methods* **2016,** 13, 875.

7.	Au - Seibenhener, M. L.; Au - Wooten, M. C. Use of the Open Field Maze to Measure Locomotor and Anxiety-like Behavior in Mice. *JoVE* **2015**, (96), e52434.

8.	Kraeuter, A.-K.; Guest, P. C.; Sarnyai, Z., The Open Field Test for Measuring Locomotor Activity and Anxiety-Like Behavior. In *Pre-Clinical Models: Techniques and Protocols*, Guest, P. C., Ed. Springer New York: New York, NY, 2019; pp 99-103.

9.	Filipe, V. M.; Pereira, J. E.; Costa, L. M.; Maurício, A. C.; Couto, P. A.; Melo-Pinto, P.; Varejão, A. S. P. Effect of skin movement on the analysis of hindlimb kinematics during treadmill locomotion in rats. *Journal of Neuroscience Methods* **2006,** 153, (1), 55-61.

10.	Au - Metz, G. A.; Au - Whishaw, I. Q. The Ladder Rung Walking Task: A Scoring System and its Practical Application. *JoVE* **2009**, (28), e1204.

11.	Minev, I. R.; Musienko, P.; Hirsch, A.; Barraud, Q.; Wenger, N.; Moraud, E. M.; Gandar, J.; Capogrosso, M.; Milekovic, T.; Asboth, L.; Torres, R. F.; Vachicouras, N.; Liu, Q.; Pavlova, N.; Duis, S.; Larmagnac, A.; Vörös, J.; Micera, S.; Suo, Z.; Courtine, G.; Lacour, S. P. Electronic dura mater for long-term multimodal neural interfaces. *Science* **2015,** 347, (6218), 159-163.

12.	Kennedy, H. S.; Puth, F.; Van Hoy, M.; Le Pichon, C. A method for removing the brain and spinal cord as one unit from adult mice and rats. *Lab Animal* **2011,** 40, (2), 53-57.

13.	Yang, X.; Zhou, T.; Zwang, T. J.; Hong, G.; Zhao, Y.; Viveros, R. D.; Fu, T.-M.; Gao, T.; Lieber, C. M. Bioinspired neuron-like electronics. *Nature Materials* **2019,** 18, (5), 510-517.
